# The 3D nuclear position and compartmentalization of genes prime their response to mechano-confinement

**DOI:** 10.1101/2025.11.06.686901

**Authors:** Saleh Oshaghi, Oda Hovet, Andrea Halaburkova, Cinzia Progida, Jonas Paulsen

## Abstract

Cells continuously receive mechanical inputs from their environment. For example, during migration in narrow spaces, or in solid tumors, cells and their nuclei experience confinement which they sense and respond to. Emerging evidence identifies the genome as a central mediator of these responses, dynamically reorganizing its three-dimensional (3D) structure to regulate both short– and long-term gene expression programs, enabling cellular adaptation to mechanical constraints. However, the mechanisms underlying such responses, especially those linking the 3D genome and transcriptome, are unknown. Here, we utilize controlled cell confinement followed by Hi-C and transcriptome analyses to map and model temporal responses to mechano-confinement in the 3D genome. We identify clusters of genes (termed “TECs”) exhibiting coordinated temporal transcriptional responses tied to their differential radial positioning within the nucleus. Additionally, we uncover a genome-wide, partially reversible response, wherein chromosomes reposition under confinement to enhance select genome compartments, aligning with temporal gene regulation patterns. Specifically, one such cluster, TEC7, is linked to 3D genome restructuring and nuclear translocation of NF-κB, driving cytokine expression and secretion. Our findings reveal how 3D genome reorganization and transcriptional programs can drive mechanoresponses, opening up for new views into the mechanisms of mechanogenomic regulation in health and disease.

## Introduction

Cells employ a range of mechanisms to sense, respond, and adapt to mechanical cues^1–6^. Increasing evidence suggests that external forces can be directly transmitted to the nucleus affecting the three-dimensional (3D) organization of the genome to impact transcriptional regulation^7,8^. This process involves force transmission from focal adhesions through actin filaments to the LINC complex, which connects to the nuclear lamina, thereby rapidly modulating nuclear morphology, chromatin structure and chromatin-lamina interactions^9,10^. The nucleus can also directly translate its shape changes into lasting mechano responses^11,12^. How such modulations trigger gene expression changes is unclear, but could happen via rapid activation of poised genes and/or determined by the spatial repositioning of genes within specific compartments^13^. Recently, we have shown that mechanical confinement creates a response in condensates of Heterochromatin 1 alpha (HP1a) by rapidly altering in number and nuclear localization^14^, suggesting that chromatin modulation is an underlying direct response to mechano confinement. Similarly, mechanosensitive nucleocytoplasmic influx of factors like HDAC3 can modulate chromatin structure and epigenetic marks to regulate gene transcription^15^.

Gene expression is influenced by the genes’ spatial positioning within the nucleus. Genes near the nuclear periphery and lamina tend to be repressed, whereas genes in the center tend to be expressed^16^. Mechano-triggered changes in gene clustering could also affect their expression^17^, suggesting repositioning within the 3D genome could affect gene regulation as a response to mechanical inputs^8^.

The 3D genome is organized into A and B compartments, representing transcriptionally permissive and repressive regions, respectively^18^. These can restructure to orchestrate gene regulation during cell differentiation^19–22^, or as a response to external stimuli^23^. Further subdivision into subcompartments reveal a gradual transition from highly permissive to strongly repressive states^24,25^, corresponding to a radial chromatin organization from the nuclear center to the periphery^26^. Alteration or loss of this radial organization has been linked to diseases like cancer^27^. Yet, how the 3D genome’s compartmental and radial organization is implicated in mechanoresponses is unclear. Studies using Hi-C, which maps 3D genome organization by deep sequencing of crosslinked and proximity-ligated DNA, have shown that constricted migration caused limited alterations in the 3D genome, despite differential expression of hundreds to thousands of genes^23^. Conversely, other studies using multiple rounds of constriction passages showed more dramatic 3D genome effects, particularly affecting B compartments^28^. These variations could be attributed to the highly dynamic and time dependent nature of chromatin mechanoresponses^29,30^. Overall, the time-dependent interplay between 3D genome organization, transcriptional responses, and mechano-signaling pathways remains poorly understood.

Here, we combine RNA-seq and Hi-C analyses to study transcriptome and 3D genome dynamics before and during a time course following mechano-confinement. Our findings reveal highly dynamic, time-dependent transcriptional responses, classified into eight distinct temporal expression clusters (TEC0 to TEC7). Intriguingly, Hi-C data show that each TEC is associated with unique subcompartment profiles, while 3D genome modeling demonstrates distinct radial positioning for each TEC. This radial organization predisposes TECs to specific functional mechano-responses, linked to their positional and subcompartment context. Moreover, we show that the 3D genome rapidly responds to mechano-confinement by a contraction of homotypic subtelomeric regions, followed by a reversal after 4 hours of confinement. We further reveal mechano-triggered nuclear translocation of NF-κB, activating a pathway that stimulates shifts in subcompartments and transcription, cytokine production and secretion, consistent with TEC7’s gene functional enrichment. Taken together, our work demonstrates how the 3D genome, transcriptome and signaling pathways combine to generate temporal mechanoresponses.

## Results

### Cell mechano-confinement reveals clusters of temporal gene-expression responses

To characterize transcriptional consequences of nuclear mechano-confinement, we performed RNA-seq of non-confined (NC) IMR90 cells, and in cells confined for either 15 minutes (C15m), 1 hour (C1h), 4 hours (C4h) or 24 hours (C24h) (see Fig. 1A-B). Inspecting temporal expression patterns during confinement, we noticed genes which were seemingly downregulated at C15m, that regained expression at C4h (Fig. 1B [left dotted box]). We also noticed genes that were turned on and gradually upregulated, maintaining their expression in C24h (Fig. 1B [right dotted box]). Other genes were non-responsive (Fig. 1B [middle dotted box]). The existence of such complex, dynamic transcriptional responses prompted us to engage a gene clustering strategy of the temporal expression patterns, using silhouette scores and hierarchical cluster analysis (see Methods). Based on logL fold-change (LFC) patterns over the course of confinement relative to NC, all genes were grouped into eight distinct clusters which we term Temporal Expression Clusters (TECs) (Fig. 1C). TEC0, which captures genes with irregular or non-responsive expression patterns, was the largest cluster with 6042 genes. TEC7, which harbors genes that are slightly downregulated at C15m, but then stably upregulated at subsequent timepoints, was the smallest cluster with 153 genes (Suppl. Table S1-S2).

**Fig. 1:**
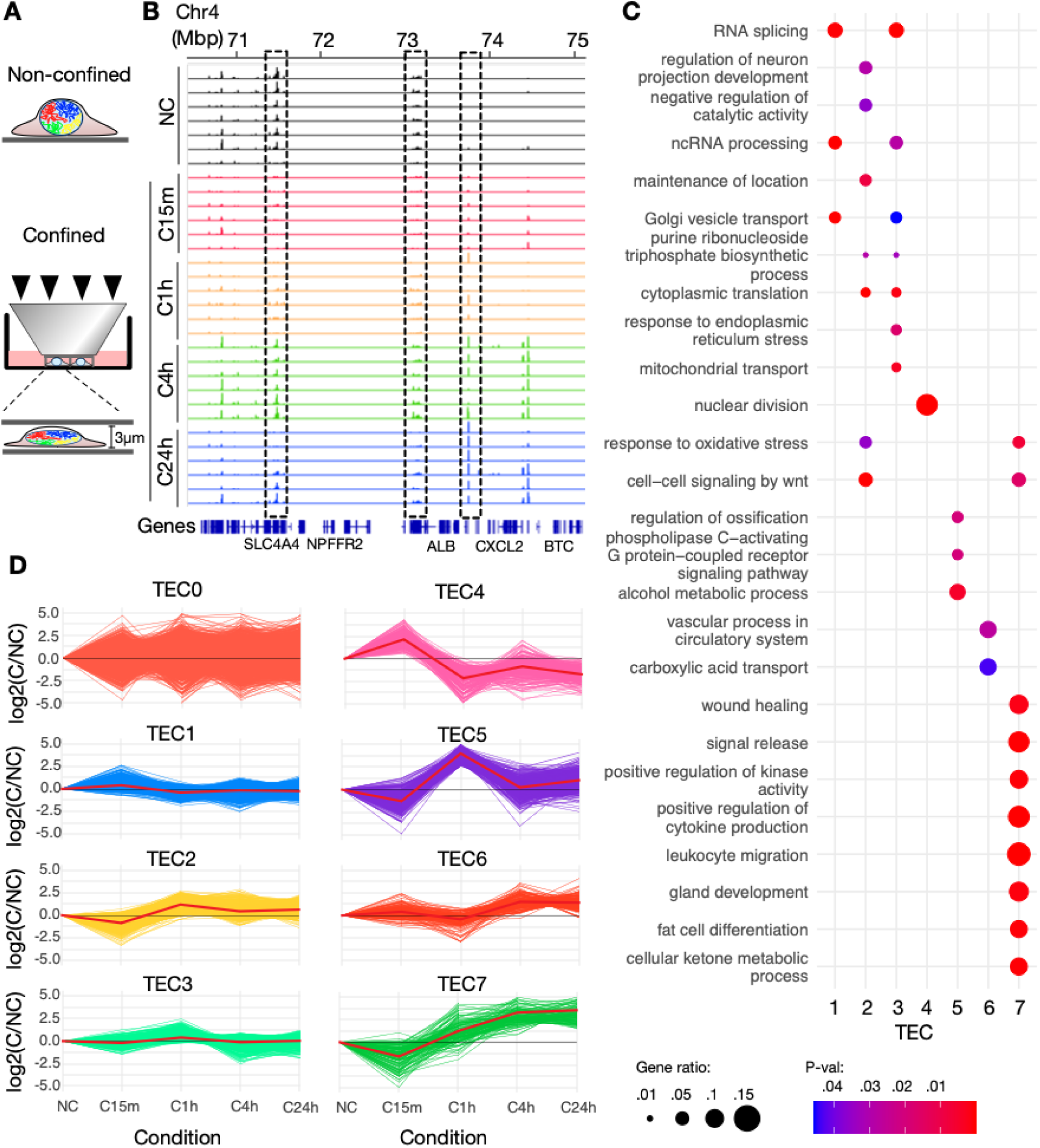
**A**: Micrometer precision static confinement system for cells and their nuclei, enabling comparative analysis of non-confined (NC) versus confined (C) cells over varying durations. **B**: RNA-seq genome tracks in an example region on chromosome 4 showing NC (black), C15m (red), C1h (orange), C4h (green) and C24h (blue). Dotted boxes highlight exemplary temporal expression patterns, with left: a gene expressed at NC, downregulated at C15m-C1h, and then upregulated. Middle: a non-responsive gene. Right: a gene gradually upregulated upon confinement. **B:** logL fold-change patterns over the course of confinement in the eight identified TECs. The red line indicates the mean value in each condition. **C:** Summarized significant Gene Ontology biological processes (BP) in each TEC. The adjusted p-value indicates the statistical significance of each BP, while the gene ratio represents the proportion of genes involved in that BP.

We wondered whether the clustering of gene expression responses into TECs could indicate that genes with similar functions responded similarly to mechano-confinement. To test this, we employed Gene Ontology (GO) analysis of each TEC separately. The resulting significant, semantically aggregated^31^ GO terms indeed revealed that each TEC harbors genes with functions that are often specific to one or a few of the TECs (Fig. 1D; Suppl. File S1). For example, TEC2 was significantly associated with the Wnt signaling pathway, with genes in this group showing, relative to NC, slight downregulation at C15m (mean LFC=-0.851; Suppl. Table S3) and mild upregulation at later time points (mean LFC=1.184 [C1h], 0.443 [C4h], 0.63 [C24h]; Suppl. Table S3). TEC4, which is upregulated at C15m and downregulated past C1h, exclusively exhibited a strong association with nuclear division and cell cycle regulation. In contrast, TEC7 was linked to the regulation of cytokine production and inflammatory response, with genes in this group initially downregulated (mean LFC= – 1.574 at C15m; Suppl. Table S3) but showing marked upregulation at C4h and C24h (mean LFC > 3; Suppl. Table S3).

To conclude, a large number of genes exhibit temporal transcriptional responses to mechano confinement which are coordinated in clusters with similar functions.

### Three-dimensional (3D) genome structure alters in concordance with TECs

We have previously shown that mechano confinement induces a flattening of the nucleus with resulting alterations in heterochromatin properties^14^. From this, we hypothesized that the three-dimensional (3D) structure of the genome could undergo a restructuring which may underlie our observed transcriptional response patterns. We noticed that many TECs show an immediate gene expression response at C15m, which often shifts in the opposite direction at C4h establishing a stable expression response seen also at C24h. To characterize the corresponding 3D genome dynamics, we generated triplicate Hi-C datasets from NC, C15m and C4h IMR90 cells (Fig. 2A; Suppl. Table S4). To facilitate a comparative 3D genome analysis, we utilized Calder to compute 8 subcompartments (Fig. 2B: A0-3 and B0-3). Overlaying the subcompartments with a range of publicly available transcription factor and histone modification ChIP-seq datasets revealed that A1-A3 subcompartments are gradually enriched in active marks including H3K9ac, H4K20me1, H3K27ac and transcription factor binding (Fig. 2C). Subcompartments B1-B3, on the other hand, show a gradual enrichment of repressive marks, including H3K9me3, LADs and NADs, which indicate constitutive heterochromatin. Notably, A0 and B0 are enriched in H3K27me3, indicative of a distinct facultative heterochromatin state, which has previously been reported in these cells^25^. Gene transcription levels are lowest for B3, and then gradually increasing towards A3, thus validating that the subcompartment assignments reflect different levels of chromatin activity.

**Fig. 2:**
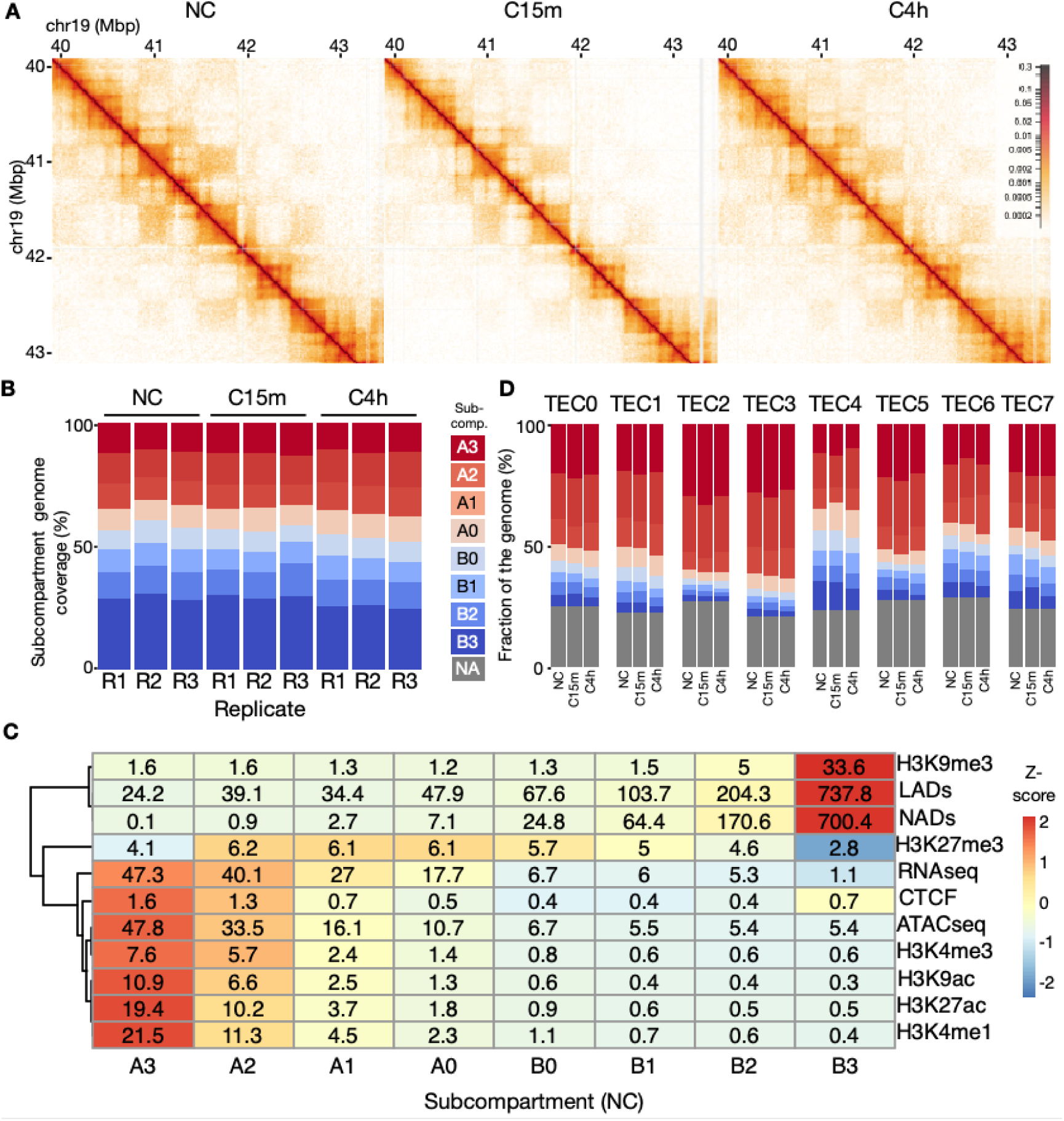
**A**: Example Hi-C region on Chromosome 19. **B:** Subcompartment composition (%) in each replicate and condition. **C:** Heatmap showing histone modification and transcription factor coverage (MBp) overlapping each subcompartment in the NC Hi-C data. The cell coloring reflects the row-scaled Z-score of each subcompartment with transcription factors, histone markers, nucleolus-associated domains (NADs), and lamina-associated domains (LADs). The RNA-seq row shows mean gene expression levels in transcripts per million (TPM) for each subcompartment. A more intense red indicates stronger association or higher expression, whereas a more intense blue indicates weaker association or lower expression. **D:** Subcompartment composition (%) by TECs determined by subcompartment state at TSS regions of each gene in each TEC. Legend on the left shows subcompartments, with gray (NA) indicating regions not assigned to a particular subcompartment (see Methods).

Overall, the genome coverage of the various subcompartments was consistent within the triplicates, and did not majorly change comparing NC, C15m and C4h (Fig. 2B). In all three conditions, B3 was the most prominent (703.8-876.4Mbp, 23.22-28.91%; Suppl. Table S5-S6), and A0 and B0 were the least prominent (188.66-288.23Mbp, 6.22-9.51%; Suppl. Table S5-S6) subcompartment. Thus, no major differences in overall subcompartment coverage were seen comparing non-confined and confined samples. We also computed Topologically Associated Domains (TADs) in all samples, which revealed an average of 3763 TADs, with minor differences between conditions (Suppl. Fig. S1-S2).

We then asked whether subcompartment associations could vary across TECs, given their distinct transcriptional profiles. To this end, we mapped subcompartment annotations onto the transcription start sites (TSSs) of individual genes within each TEC. Our analysis revealed that each TEC indeed exhibited a distinct subcompartment composition (Fig 2D). For example, TEC4, which harbors genes associated with nuclear division (see Fig. 1C) which become stably downregulated at C1-24h, displayed an enrichment of constitutive heterochromatin subcompartments (B2-B3) alongside a higher fraction of facultative heterochromatin (A0-B1) compared to other TECs. In contrast, TEC2, with a transient downregulation at C15m and modest upregulation at later time points, showed a relative enrichment of A3 subcompartments at the expense of A0-B3. Overall, TECs display a higher fraction of A2-A3 subcompartments compared to the genome average (see Fig. 2B), which is not surprising given that gene regions are generally found in more active parts of the genome. In conclusion, genes within TECs harbor distinct subcompartment profiles at their TSSs.

Because subcompartment association differed between TECs, we wanted to assess whether TECs were also differentially positioned in the 3D genome. To this end, we generated whole genome 3D models from the Hi-C data using Chrom3D^32^. In these models, each TAD is modeled as “beads-on-a-string” per chromosome (Fig. 3A), which is optimized based on significantly interacting regions identified from Hi-C data. Overlaying subcompartments onto each bead in the resulting Chrom3D models (Fig. 3B) revealed that subcompartments from A3 to B3, gradually position from central to peripheral nuclear regions, respectively (Fig. 3C). For example, in NC, A3 is located on average 3.35 μm from the nuclear center, compared to B3 at 3.75 μm (Suppl. Table S7). This difference is statistically significant based on a Wilcoxon signed-rank test (P = 7.12e-15; Suppl. Table S8). This trend is also seen in C15m and C4h (Suppl. Table S9-S10), and confirms that our computed subcompartments reflect expected radial chromatin organization^26^, and also validates our Chrom3D models.

**Fig 3:**
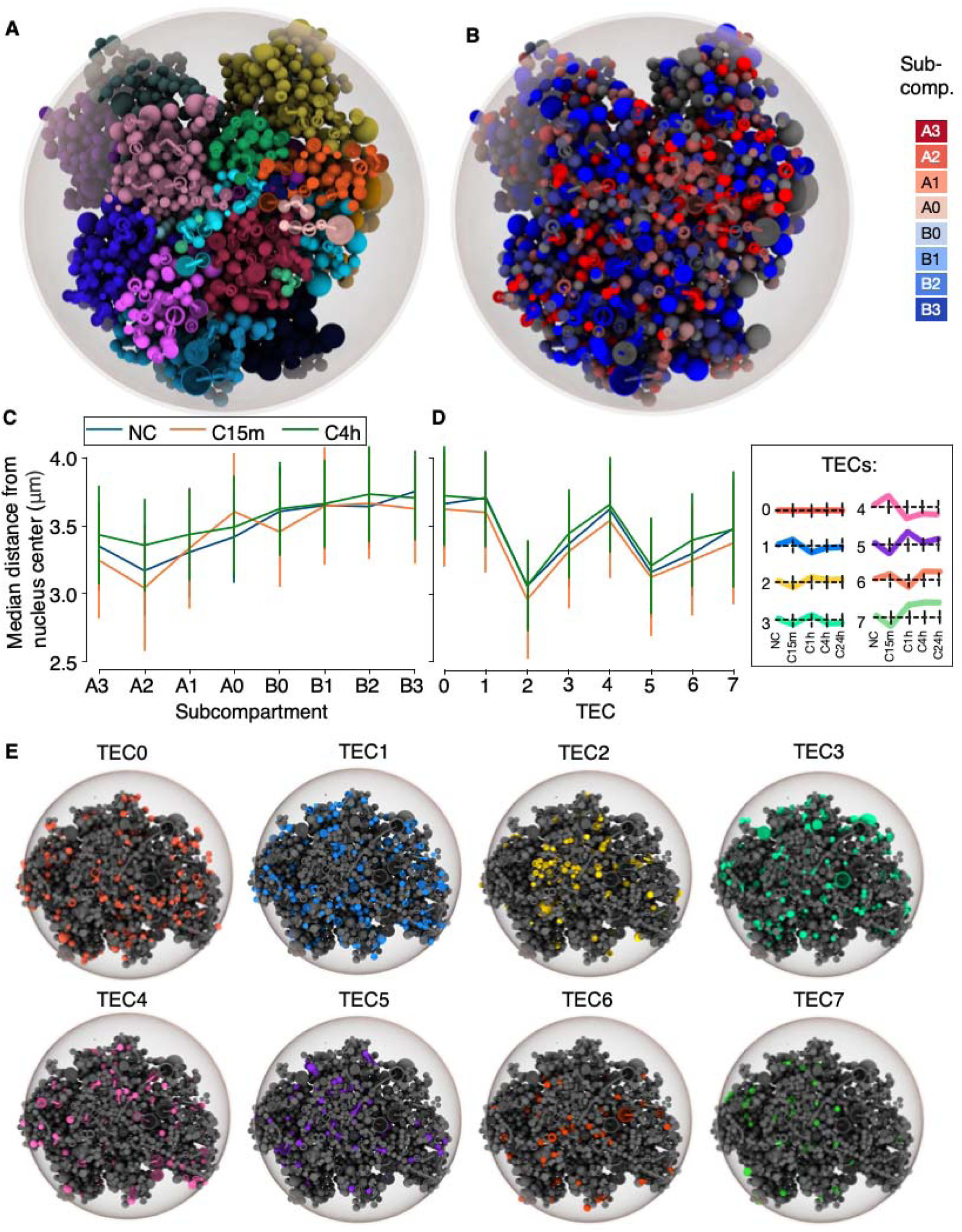
**A**: Left: Example tomographic view of a NC Chrom3D model with chromosomes colored individually. **B:** The same model, with regions color-coded according to their subcompartment assignments. **C:** Median ± standard deviation of distances from the nuclear center for each subcompartment under conditions NC, C15m, and C4h. **D:** Median ± standard deviation of distances from the nuclear center for each TEC under conditions NC, C15m, and C4h. **E:** Exemplary Chrom3D models (C15m) rendering all genes in each TEC.

Performing a similar radial comparison of the subcompartments contrasting NC, C15m and C4h, revealed that most subcompartments remain relatively stably positioned upon confinement (Fig. 3C; Supp. Table S11). The most notable change was seen for A0 subcompartments, which compared to NC, peripheralize from 3.42μm to 3.60μm at C15m (Suppl. Table S7; P=2.20×10^-7^ [Suppl. Table S11]), but partially centralize again to 3.49μm in C4h (Suppl. Table S7; P=0.003 [Suppl. Table S11]). This trend seems specific to A0 subcompartments, which notably shifts its association with facultative heterochromatin marks (i.e. H3K27me3) in NC compared to C15m and C4h (Suppl. Fig. S3). This suggests that subtle changes in facultative heterochromatin also could underlie genomic responses to confinement.

In line with their differential association with subcompartments at gene TSSs, we wondered whether different TECs could be positioned in distinct nuclear regions. We thus overlapped TSSs of genes in TECs onto the TADs making up our Chrom3D models, revealing a striking difference in their radial positioning (Fig. 3D). For example, genes in TEC2 were across conditions placed towards the nuclear center (2.96-3.06μm; Suppl. Table S12), whereas genes in TEC4 were placed towards the nuclear periphery (3.54-3.66μm; Suppl. Table S12). Using a Wilcoxon rank-sum test, this difference in radial positioning is statistically significant (P<8.17e-18; Suppl. Table S13-15) and clearly visible when rendered on individual Chrom3D models (Fig. 3E).

When analysing differences in radial positioning of TECs between conditions (NC, C15m and C4), radial shifts are modest and generally less than +/-0.1μm (Suppl. table S16). The largest radial difference is seen for TEC3, which shifts from 3.31μm (C15m) to 3.44μm (C4h) (Suppl. Table S12; P=0.003 [Suppl. Table S16]). Across TECs, all genes shift slightly towards the nuclear center at C15 (median shift –0.02μm; P=1.43e-06; Suppl. Fig.S4), and revert to a slightly more peripheral position at C4h (+0.01μm; P=0.000178; Suppl. Fig. S4). Thus, whereas the radial positioning of the different TECs varies greatly, their positions remain rather stable upon confinement, suggesting that TECs are spatially prepatterned to engage specific temporal transcriptional programs upon external stimuli.

We then asked whether the observed differences in subcompartments and nuclear radial placement between TECs is indicative of a tendency for subcompartments to switch as a response to mechano-confinement. Genome-wide, contrasting NC and C15m (Suppl. Fig. S5) and C4h (Suppl. Fig. S6) reveals a modest degree of switching mainly involving consecutive subcompartments. When contrasting switching between TECs only (Suppl. Table S1), the fraction of genes not switching between NC and C15m varies moderately from 42% (TEC6) to 48% (TECs 3,4 and 7). Similarly, contrasting NC and C4h reveals a fraction of non-switching genes varying from 32% (TEC6) to 43% (TEC3). Classifying switching into “chromatin opening”, which is when a subcompartment changes toward a more open or A-like subcompartment, or vice versa for “closing”, reveals that TEC4 has a slightly lower fraction of chromatin opening (14%, whereas remaining TECs have 16-18%) and higher fraction of chromatin closing (15%, compared to 9-14%) contrasting NC and C15m. This is in line with TEC4’s downregulation at 1h of confinement (see Fig. 1B), and its association with heterochromatic subcompartments (Fig. 2D) and the placement of its genes toward the nuclear periphery. Generally, however, TECs display a similar degree of switching to opening and closing chromatin. Similarly, classifying switching into a gain or loss of facultative subcompartment types (A1-B1) reveals no striking differences in types of switching between TECs contrasting either NC and C15m (Suppl. Table S1) or NC and C4h (Suppl. Table S2).

Because the direction of the differential expression of the TECs often was opposite in C15m compared to C4h, yet tended to stabilize after C4h, we classified TECs based on their upregulation (TECs 2, 5, 6, 7) or downregulation at (TECs 1, 3, 4) in C4h relative to NC. With these two groups (i.e. upregulated and downregulated TECs), we wanted to analyze whether their genes harbor differential switching of subcompartments, and used a Fisher’s exact test to analyze subcompartment opening and closing at their TSSs. At C15m, subcompartment opening and closing were significantly linked to up-or downregulation, respectively (P=3.805e-05; Odds ratio = 0.723; Suppl. Table S17), but the trend reversed at C4h (P=0.0008173; Odds ratio = 1.27386). This suggests that gene expression patterns at C4h and onwards are determined immediately (i.e. within 15 min) to impact gene expression patterns responsive to confinement (Suppl. Table S17).

In conclusion, TECs are placed in distinct 3D genome regions within the nucleus, with a tendency for more active subcompartments and a positioning towards the nuclear center for TECs displaying upregulation after confinement (TEC2 and 5-7), and more inactive subcompartment and peripheral nuclear placement for TECs either nonresponsive (TEC0) or downregulated after confinement (TEC1, 3 and 4). Thus, the 3D genome positioning of TECs in non-confined cells could prepattern genes towards specific transcriptional responses to nuclear confinement.

### The 3D genome rapidly responds to mechanical inputs in a semi-reversible manner

Beyond gene-centered responses to mechano-confinement, we hypothesized that a confinement-induced flattening of the nucleus also should reorder the global 3D genome potentially driving responses to mechano-inputs. To analyze such genome-wide reorganization, we employed a multi-dimensional scaling (MDS) method on all interchromosomal Hi-C contact frequencies between all somatic chromosomes in NC, C15m and C4h to optimize the relative positioning of each chromosome on a 2D plane. The resulting MDS plots (see Fig. 4A) revealed a striking chromosome positional response at C15m, wherein most chromosomes come together, followed by a reversal of their relative position at C4h. For example, chromosome 9 (Fig 4A; red color) comes together with chromosomes 8, 10-12, 15-17,19 and 22 in C15m, but restores a similar relative location as in NC in C4h (Fig. 4A; right panel).

**Fig. 4:**
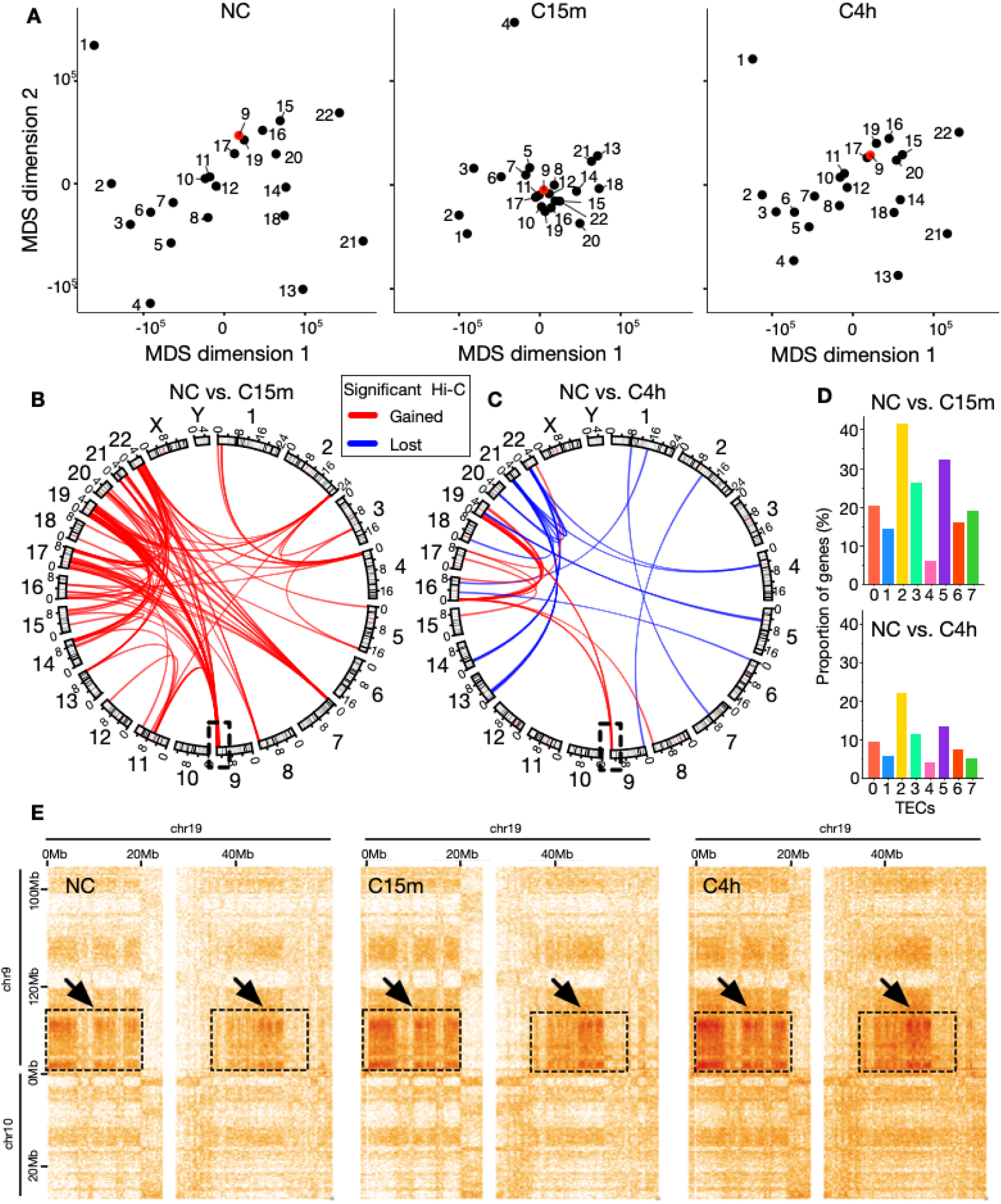
**A**: MDS plot based on the interchromosomal Hi-C contact frequencies, showing the positions of each somatic chromosome (dots) along the first two dimensions in NC (left), C15m (middle) and C4h (right). Chromosome 9 is highlighted in red. **B:** circos plot showing significantly enriched (red) and depleted (blue) inter-chromosomal contacts in NC vs. C15m. and in (**C**) NC vs. 4h. Genomic positions along angular axes are specified in 100Mbps; **D:** Proportion of genes (%) for each TEC in regions with differential interactions (see B and C) in NC vs. C15m (top panel), and NC vs. C4h (bottom panel). **E:** Example Hi-C map showing significantly enriched interactions between genomic regions on chromosome 9 (130,000,000–138,394,717 bp) and chromosome 19 (0–55,000,000 bp). Arrows pointing to dotted squares indicate regions where there are statistically significant gains of contacts.

To assess whether the confinement-induced semi-reversible chromosomal repositioning was driven by more specific regions on the chromosomes, we divided the genome into 5Mb-sized bins and analyzed statistically significant differential Hi-C contacts between all pairs of bins using Deseq2 (see methods). This analysis identified 110 significant inter-chromosomal differential contacts between NC and C15m (Fig. 4B) (out of 228 total intra– and interchromosomal differential contacts; Supplemental File S2), and 36 differential inter-chromosomal contacts contrasting NC and C4h (out of 1530 total; Supplemental File S3) (Fig. 4C). Interestingly, these differential contacts were primarily enriched in subtelomeric regions of chromosomes. They appeared exclusively gained at C15m (Fig. 4B; 110 interactions) but some were later lost at C4h (Fig. 4D; 20 lost interactions), frequently affecting many of the same chromosomes yet in distinct regions. An exception was the subtelomeric region on chromosome 9 (Fig. 4B; highlighted square), which gained interactions in the same region at both C15m and C4h.

Overlapping genes from each TEC with regions of gained and lost contacts indicated that TEC2 and TEC5, the two TECs most centrally placed in the nucleus, were more likely to overlap with gained interactions at C15m compared to NC. TEC4, the most peripheral TEC, on the other hand, had the least overlap (Fig. 4D; top). Comparing the gained and lost interactions in C4h relative to NC, revealed a similar trend, yet with lower overlap overall (Fig. 4D; bottom).

To investigate the details of the gained differential contacts, we zoomed in on the Chromosome 9 region in the Hi-C data harboring increased contact frequencies with regions on chromosome 19 (Fig. 4E; arrows). Investigating these regions further, revealed that they harbor a large fraction of A2 and A3 subcompartments (Suppl. Fig. S7), which thus specifically strengthen their contacts in response to confinement. Indeed, across all regions with gained or lost contacts as a response to confinement, the tendency is for these interactions to be within-compartment (homotypic) interactions (Suppl. Fig. S8-S9).

### Mechano-confinement induces cytokine response and release

Among the different TECs, TEC7 showed a gradually increasing expression of genes involving cytokine production, signal release, Wnt-signalling and wound healing (Fig. 1C). A known key factor driving these processes is the transcription factor NF-kB, which rapidly shuttles to the nucleus to bind regulatory elements as a response to external stimuli^33^. Immunostaining experiments using antibodies against either NF-kB or the phosphorylated serine 529 of its p65 subunit (pNF-kB) validated their involvement, as 5 minutes of confinement translocates them to the nucleus (P<2.2e-16 [Suppl.Table S18-19]; Fig 5A-B; Suppl. Fig. S10). To explore the involvement of NF-kB in relation to our TECs, we analyzed the presence of genes belonging to the NF-kB-pathway (from KEGG) across TECs, revealing that TEC7 indeed is regulating NF-kB-pathway genes^34–36^ (Fig. 4C; top). Expanding this analysis to NF-kB target-genes using known transcription factor binding site motifs^37–39^, also reveals a strong association to TEC7 (Fig. 5C; bottom).

**Fig. 5:**
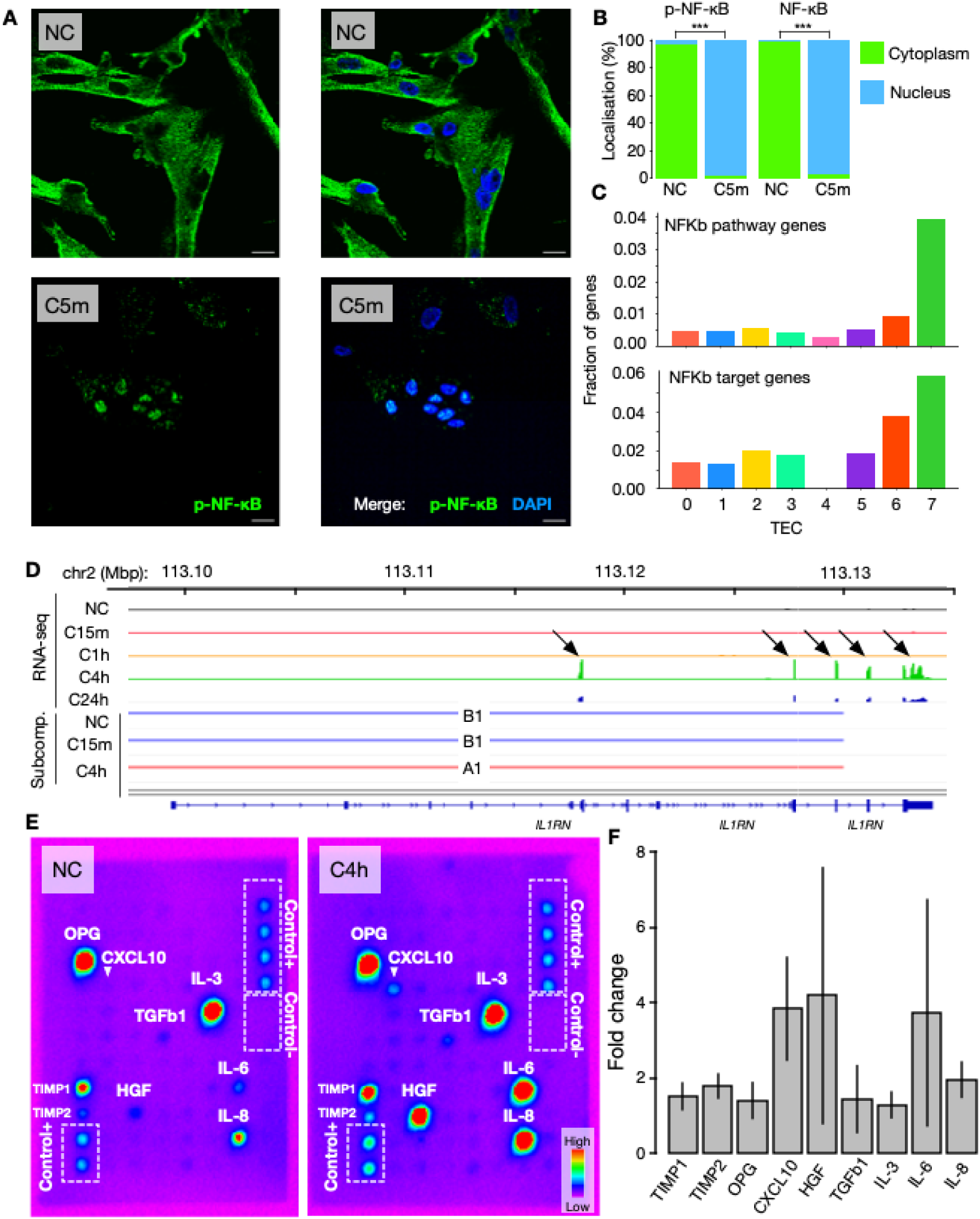
**A**: Immunostaining of p-NF-kB (green), and DAPI (blue) in NC (top) and cells confined for 5 minutes (C5m; bottom). Merge shown in the right panels. Scale bar = 20 μm. **B**: Percentage of cells with p-NF-kB or NF-kB in the cytoplasm (green) or the nucleus (blue); P<2.2e-16 (Fisher’s exact test; Suppl. Table S18-19). **C:** upper panel: NF-kB KEGG pathway gene fraction by TEC; lower panel: NF-kB target gene fraction by TEC using the Harmonizome gene target database. **D:** An example of a TEC7 gene, *IL1RN*, indicating subcompartment switch in accordance with its upregulation in C4h. The RNA-seq tracks show merged gene expression under conditions NC, C15m, C1h, C4h, and C24h. The subcompartment tracks display consensus subcompartments for conditions NC, C15m, and C4h. **E**: Cytokine antibody array on NC (left) and C4h (right) cells, showing relative signal intensities. **F**: Fold change in the signal intensity of detected secreted cytokines from the cytokine array in NC vs C4h. Values are mean ± s.d. of 2 independent experiments (Suppl. Table S20).

The observation that confinement-induced nuclear translocation of NF-kB is engaging a TEC7-coordinated gene expression response by first decreasing (after 15 minutes), and then stably increasing (after 1 hour of confinement), prompted us to analyze whether NF-kB target genes generally engage in subcompartment switching more than genes overall. To this end, we analyzed presence/absence of NF-kB target genes in (non-)switching regions revealing that at C15m indeed displays more switching of NF-kB target genes than non-target genes. (OR=1.33; P=0.022; Fisher’s exact; Suppl. Table S21). For example, the TEC7 gene *IL1RN*, an IL-1 antagonist, is lowly expressed in NC-C1h, but switches from B1 to A1 at C4h in concordance with an upregulation of its expression (Fig. 5D).

We next assessed whether the TEC7-related confinement response with increased expression of genes encoding for cytokines results in augmented cytokine secretion. We utilized a cytokine assay probing secretion of 80 different factors (Fig. 5E; Suppl. Fig. S11). From this, we observed a range of interleukins and cytokines secreted at higher levels in C4h relative to NC (Fig. 5E). Among them, 9 were visibly secreted and clearly increased, including CXCL10, HGF and IL-6 and IL-8 that showed the highest increase in secretion in confined cells (Fig. 4F; Suppl. Table S20).

To conclude, we demonstrate a mechano-induced response starting from the nuclear translocation of NF-kB, via a TEC7-associated transcriptional regulation involving switching of subcompartments leading to increased expression and secretion of cytokines.

## Discussion

We show that mechano-confinement induces temporal gene expression responses classified into eight clusters termed TECs. Further analysis reveals that each TEC exhibits i) unique gene functional enrichment, ii) distinct radial nuclear positioning, and iii) a specific subcompartment profile. Intriguingly, TECs positioned more towards the nuclear periphery, such as TEC4 (nuclear division) and TEC1 (RNA splicing; ncRNA processing; Golgi vesicle transport), are upregulated in response to 15 minutes of mechano-confinement, but then downregulated in subsequent time points. More active TECs positioned towards the nuclear center, however, such as TEC2 (Wnt signalling, among others) and TEC5 (signaling and metabolic processes) show the opposite pattern, whereby genes are first downregulated at C15m, and then upregulated in later time points. This proposes that the radial position and subcompartment profile of the TECs in non-confined cells predispose their genes for particular transcriptional responses to confinement. Furthermore, our Hi-C data and Chrom3D modeling show that all TECs respond by moving closer to the nuclear center at C15m, and then returning to their radial location at C4h (Fig. 3D), suggesting that confinement induces an immediate radial response for all TECs, followed by an incomplete reversal process. For most TECs, the original expression pattern then re-establishes with exaggerated polarity, wherein central TECs become even more active, and peripheral TECs more repressed. It may be that flattened nuclei during confinement could generate an increase in the distance of equatorial regions from the nuclear center, amplifying the radial distinction and driving more extreme gene activity differences (Fig. 6).

**Fig. 6:**
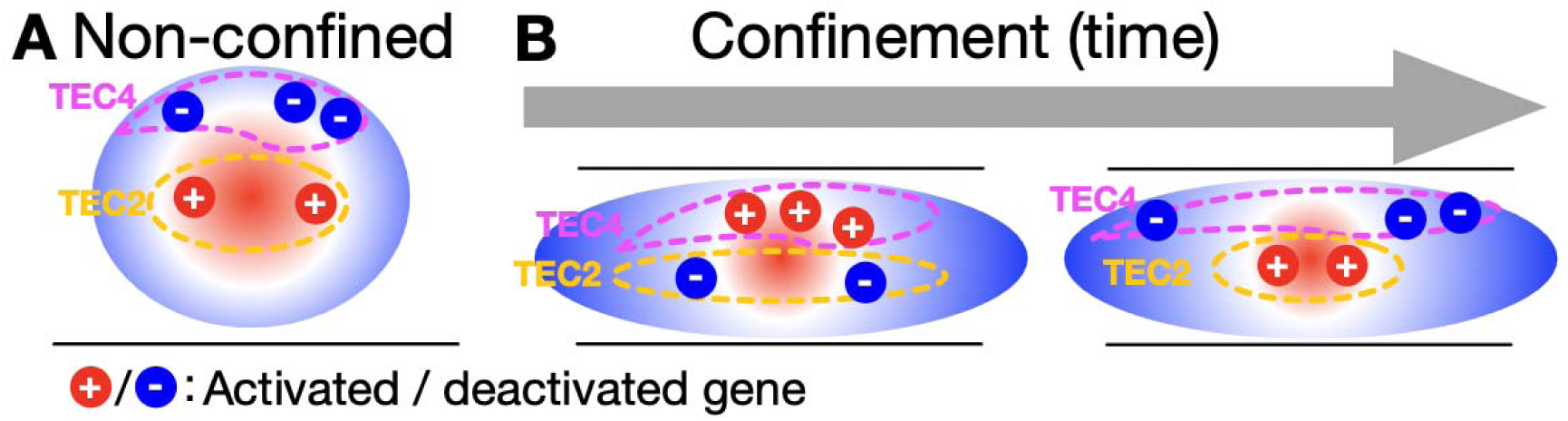
**A**: Genes (red and blue dots) in TECs (purple and yellow dotted lines) are placed in different radial compartments in the nucleus (indicated by a sphere with red and blue shading indicating A and B compartmentalization, respectively) in non-confined cells. **B**: The radial organization of the nucleus (active center, inactive periphery) and TECs’ relative placements are maintained during confinement, despite interactions within TECs being e.g. pulled closer together (e.g. at C15m). However, confinement disrupts the radial relationship between TECs/genes and the nucleus, temporarily shifting activity: peripheral TECs tend to activate, while central TECs tend to deactivate. Over time, as TEC placements are restored, the original expression pattern reestablishes with exaggerated polarity—central TECs become even more active, and peripheral TECs more repressed thus driving the temporal expression patterns of these TECs.

Intriguingly, the same confinement system we used in our study was previously shown to activate a NF-κB-dependent shape-sensing pathway in dendritic cells, triggering the production of cytokines^40^. Our data show that the same pathway is also activated in fibroblasts, suggesting that this may be a more general mechano-sensoring pathway activated by compression not only in immune cells, despite the specific set of produced cytokines may be cell dependent. Further supporting this, mouse cardiac fibroblasts activated by pressure overload highly express NF-κB target genes, chemokine and cytokine encoding genes, involved in the recruitment of monocytes to the heart^41^. Similarly, mechanical forces like compression have been reported to activate the NF-κB pathway and proinflammatory cytokine production in human periodontal ligament stem cells^42^.

Mechanoresponses in the 3D genome depend on both cell type, stimulus type and substrate, and can involve both increase or decrease in heterochromatin and other epigenome modifications^13^. These processes are increasingly acknowledged as dynamic, unfolding across multiple timescales and hierarchical levels. They encompass an integrated temporal response involving biochemical signaling pathways, chromatin and genome remodeling, and gene expression changes^30^. Our study provides a novel view into such dynamic mechanogenomic responses, which include transient and stable changes at the transcriptional and 3D genome levels. This underscores the importance of integrating temporal dynamics in mechanogenomic research for a comprehensive understanding of the genome’s response to mechanical stimuli.

## Methods

### Cells

IMR90 cells (ATCC) were grown in Eagle’s Minimum Essential Media (EMEM; Lonza) supplemented with 10% FCS, 2 mM L-glutamine, 100 U/ml penicillin, and 100 μg/ml streptomycin at 37 °C with 5% CO_2_. Cells were routinely tested for mycoplasma.

### Confinement

IMR90 cells were seeded on MatTek glass-bottom dishes. Confinement was performed using a 6-well plate confiner or 1-well plate confiner (4DCell) as we previously described in^14^. The cells and their nuclei were confined to a defined height 3-5 μm, for 5, 15 minutes, 1 hour, 4 hours or 24 hours.

### Cytokine assay

Cell culture media from NC and after C4h were collected and analyzed using the C5 cytokine antibody array (RayBiotech, AAH-CYT-5) for the semi-quantitative detection of 80 proteins, following the manufacturer’s protocol (see Suppl. Fig. S11). Signal intensities were quantified using ImageJ^43^, normalized to internal positive controls, and expressed as fold changes according to the manufacturer’s protocol, where fold change represents the relative expression of C4h compared to NC.

### Immunostaining

For immunofluorescence cells were seeded on MatTek glass-bottom dishes. NC and cells under confinement were washed with 1X PBS, fixed with 3% PFA for 20 minutes, washed with 1X PBS, quenched using 50 mM NH_4_Cl for 10 min and then de-confined. Cells were permeabilized and washed by using 0.25% Triton X-100 in 1X PBS for 5 minutes and incubated with primary antibodies for 30 minutes at room temperature. Cells were subsequently washed with 0.25% Triton X-100 in 1X PBS three times, incubated with secondary antibodies for 20 minutes at room temperature, washed again three times with

0.25% Triton X-100 in 1X PBS, and incubated with 0.1 μg/ml DAPI (Sigma-Aldrich) in 1X PBS. The samples were imaged in 1X PBS using an Olympus SpinSR SoRA spinning disk confocal microscope equipped with a PLAPON 60×/1.42NA oil objective. Z-stacks with 0.24 μm step size were generated with a resolution of 2047 × 2035 pixels. The following primary antibodies were used for the IF staining: anti-phosphoNF-kB (1:100, SC-166748, Santa Cruz) and anti-NF-kB (1:100, SC-8008, Santa Cruz).

The quantification of protein localization was based on the presence of signal in cells in the cytoplasm and/or nucleus. Cells showing signals in both were classified as cytoplasm.

### Hi-C library preparation and steps

Briefly, approximately 10^6^ cells were collected and centrifuged at 585 g for 5 minutes. Pellets were mixed by inverting 5 times with 1X PBS containing 3% BSA (v/v) and fixed with 37% formaldehyde at RT for 10 minutes with occasional inversion. Hi-C data were generated using the Arima-HiC+ kit, and the Arima Library Prep Module according to the manufacturer’s protocols. Sequencing was done by Illumina NovaSeq X paired end 150bp x 2.

### RNA-seq library preparation and steps

Total RNA was extracted using the RNeasy Mini kit (Qiagen). RNA-seq data was generated using the TruSeq mRNA-seq library (Illumina) following the manufacturer’s protocol and sequenced on NovaSeq X in two batches. The RNA-seq experiment included seven replicates for NC and six replicates for each of the confined conditions C15m, C1h, C4h and C24h.

### Hi-C preprocessing

The human reference genome (GRCh38/hg38, version p14) was obtained from the UCSC Genome Browser database^44^. Samtools was used to filter the FASTA file from mitochondrial chromosomes and unplaced scaffolds^45^. The nextflow nf-core/hic version 2.1.0 was used to perform Hi-C raw data analysis including quality control, mapping and detecting valid interaction pairs^46–53^. The restriction sites ^GATC,G^ANTC,C^TNAG, and T^TAA and their corresponding ligation sites were specified. The Bowtie2 aligner^53^ was run with parameters ‘––trim5 5 ––very-sensitive –L 30 ––score-min L,-0.6,-0.2 ––end-to-end ––reorder’, including a 5bp trimming at the 5’ end. Valid Hi-C pairs were loaded into Cooler to generate contact matrices at 1 kbp resolution^54,55^. The Cooler merge, zoomify and balance commands were used to merge the triplicates for each condition, create a .mcool file with multiple resolutions, and normalize the Hi-C matrix using standard parameters. All commands used are provided in the hic_preprocessing.txt file in our Git repository.

### RNA-seq preprocessing and differential expression analysis

RNA-seq preprocessing was performed using the Nextflow v22.10.7 nf-core/rnaseq pipeline v3.10.1^46^. Read quality control was processed by FastQC v0.11.9^56^. Subsequently, TRImgalore software v0.6.7^57^ was used to remove adapter sequences and low-quality bases. The FASTA file for reference genome hg38.p14^58^ was filtered with SeqKit v2.5.1^59^ to remove mitochondrial chromosomes and unplaced scaffolds with the pattern ‘^chr[XY\d]+$’. The reads were aligned to this reference genome using STAR v2.7.9a^60^. Gene-level counts were generated from the alignments with featureCounts v2.0.1^61^ using the Gencode v43 genome annotation^62^. Transcript-level abundance was estimated using the pseudo-aligner Salmon v1.9.0^63^. The raw gene counts table was further used for downstream analysis in R. The command for running the RNA-seq preprocessing steps is included in rnaseq_preprocessing.txt in our Git repository.

Differential expression analysis was performed on the raw gene counts table using the DESeq2 R package v1.38.3^64^. One sample from the C15m condition (15m_REP2) was excluded from this downstream analysis due to substantial deviation from other replicates within the same condition as revealed by Principal component analysis (PCA) plots (see Suppl. Fig. S12A-B). The differential expression analysis compared the time points of confinement (15m, 1h, 4h and 24h) to the control group (NC) taking the conditions and batch effects into consideration in the design parameter. Genes with an adjusted p-value < 0.05 were considered significantly differentially expressed. The code is provided in the diff_exp_analysis.R file in our Git repository.

### Temporal expression clustering and gene ontology analysis

The log2FoldChange values of NC vs. C15m, C1h, C4h, and C24h were obtained in the differential expression analysis (see *RNA-seq preprocessing and differential expression analysis*) and imported into R (R Core Team, 2025) using the readxl package. Genes with undefined log2FC values were discarded, resulting in 17,254 genes with valid log2FC values across all conditions. The resulting table was used to perform hierarchical clustering with the agglomeration method *ward.D2* using the Cluster package^65^. The dynamicTreeCut package^66^ was applied to the hierarchical clustering dendrograms for adaptive branch pruning with a minimum cluster size of 100 and a deepSplit value of 0. Subsequently, silhouette information was computed using the cluster package, and genes with a silhouette value below 0.1 were assigned to TEC0, indicating irregular expression patterns (Suppl. File S4). The code is provided in the TEC_clustering.R file in our Git repository.

The list of NF-κB pathway genes was obtained from the KEGG database^34–36^, and the list of NF-κB target genes was obtained from the Harmonizome 3.0 database^37,38,67^, which is based on known transcription factor binding site motifs. These lists were intersected with the set of genes annotated to clusters using the Python packages bioframe^68^ and pandas^69^, and visualized with matplotlib^70^.

Gene symbols were converted to Entrez IDs using the R package *AnnotationDbi* together with the database *org.Hs.eg.db*. Gene ontology analysis was performed with *clusterProfiler*^71^ using the function *enrichGO* with the biological process (“BP”) category, a p-value cutoff of 0.05, and the Benjamini–Hochberg (“BH”) method for multiple testing correction. The results were exported with the *rio* package.

The top 50 most significant GO terms with at least 10 genes per TEC were summarized using semantic similarity measures calculated with the R package *GOSemSim*^31^. A similarity matrix was generated, and GO terms with a minimum semantic similarity score of 0.3 were grouped into interconnected components using the *igraph* package. Within each group, the most significant GO term was selected as the representative (Figure 1C). The code is provided in the TEC_GO_analysis.R file in our Git repository.

The R packages dplyr, tidyr, tidyverse, stringr, ggplot2, ggforce, gridExtra, and cowplot and the Python packages numpy, bioframe, pandas, and matplotlib were used for data transformation and visualization. The heatmaps were created in R using pheatmap. Figures from the Hi-C matrices were created using HiGlass^72^, and genome browser views were generated with IGV^73^.

### Subcompartment analyses

CALDER2^74^ was employed for subcompartment calling and installed directly from the CSOgroup/CALDER2.0 GitHub repository using the remotes package, which was installed via BiocManager. Following CALDER2 recommendations, GenomicRanges^75^ was installed separately. As CALDER2 requires .hic files, .cool files were converted using hictk^76^. The resulting .hic matrices were normalized using hictk balance with ICE (iterative correction and eigenvector decomposition) and the parameters ––no-rescale-weights and ––mode=cis. Subcompartment analysis with CALDER2 was performed at 5 kb resolution on the hg38 genome assembly, without employing the subdomains functionality (Code: CALDER2_subcompartment_calling.R). For each condition, a consensus subcompartment was assigned to regions exhibiting the same subcompartment in at least two of the three replicates.

A heatmap showing subcompartment switches in NC vs. C15m and NC vs. C4h was generated using the pheatmap package to visualize the degree of subcompartment changes in Mbp (Supplementary Figures S5 and S6; code available in *subcom_viz.R*). Fisher’s exact test was used to assess subcompartment switches in NF-κB target genes compared with non-NF-κB target genes, implemented with the fisher_exact function from the SciPy package^77^. Fisher’s exact test in base R was used to statistically evaluate chromatin opening versus closing in the upregulated clusters (TEC 2, 5, 6, and 7) compared to the downregulated TECs (TEC 1, 3, and 4).

Publicly available data were retrieved from the ENCODE portal (https://www.encodeproject.org/)^78^ for subcompartment overlap comparison analyses. A complete list of the datasets used, along with their respective accession numbers, is provided in Supplementary Table S22. UCSC LiftOver was applied to convert genomic coordinates to the GRCh38 assembly when GRCh38 coordinates were not available^79^. These domains were intersected with subcompartment domains using Bioframe, and the log2-transformed total overlap length is shown in Figure 2C.

### 3D genome modeling

The function hicFindTADs from HiCExplorer v3.7.4^80^ was used to identify topologically associating domains (TADs) for all replicates and conditions at 50 kbp resolution with “–– correctForMultipleTesting fdr,” using Hi-C files normalized with the script cooler_apply_normalization.py available at https://github.com/paulsengroup/2022-mcf10a-cancer-progression. Boxplots of the total number of length of the TADs were created in R using ggplot2^81^ (Suppl. Fig. S1 and S2). Consensus TAD boundaries were defined by comparing all boundaries across samples and selecting those present in at least 5 of 9 samples, ensuring consistent chromatin domains for subsequent 3D genome modelling and comparative analysis.

The NCHG package (https://github.com/paulsengroup/NCHG.git)^82^ was used to detect significantly interacting chromatin domains. Using the *NCHG cartesian-product* function, we generated a BEDPE file representing the whole genome segmented by the consensus TAD domains. Expected contact matrices were computed with *NCHG expected* at 50 kbp (cis) and 1 Mbp (trans) resolutions. These matrices were then used in *NCHG compute* to identify significant interactions, followed by the *NCHG merge*, *filter*, and *view* steps to extract statistically significant interactions (FDR = 0.01; logRatio = 1.5 for interchromosomal and 2.0 for intrachromosomal interactions). The total number of significant interactions per replicate is reported in Supplementary Table S23. Only inter– and intrachromosomal interactions that were significant in at least two of three replicates per condition were retained for downstream 3D genome modelling.

The statistically significant chromosomal interactions were converted into gtrack files using the hg38 genome size file and the TAD segmented genome. A patched version of Chrom3D v1.0.2^32^ was then used to generate 100 Chrom3D models for each condition (NC, C15m, C4h) at TAD resolution, with a nuclear occupancy parameter of 0.15 and 2 × 10L simulation steps (Code at chrom3d_modeling.sh). ChimeraX^83^ was used to visualize the simulated Chrom3D models and to generate all Chrom3D figures included in this manuscript.

### Chrom3D downstream analysis

The subcompartments were mapped onto the beads in the Chrom3D models, with each bead assigned to the subcompartment that had the largest overlap. Similarly, gene coordinates were mapped onto the beads, and each bead was assigned to the temporal expression cluster (TEC) that contained the largest proportion of its associated genes. The median distance from the nucleus center and its standard deviation for each subcompartment and TEC were then calculated and visualized. A pairwise Wilcoxon rank-sum test was performed between all subcompartments and clusters in each condition (NC, C15m, and C4h) using pairwise.wilcox.test in R (Suppl. Tables S8-S11 and S13-16). For each bead in the Chrom3D models corresponding to a TEC, the median distance from the nuclear center was calculated. The distribution of differences in median distances between NC vs. C15m and NC vs. C4h was visualized as a histogram (Supplementary Fig. S4). Pairwise comparisons of the median distances between NC vs. C15m and NC vs. C4h were performed using the Wilcoxon rank-sum test implemented in the wilcox.test function in R.

### 3D genome reorganization

To ensure a fair comparison of chromatin interaction counts between the different conditions (NC, C15m, and C4h), the merged .mcool files for NC and C4h were downsampled to match the total sum of the C15m .mcool file, which had a total sum of 1,205,642,737. This was done using cooltools random-sample^84^. The downsampled .mcool files were then balanced and zoomified using cooler^54^.

The average balanced interaction frequency between each pair of chromosomes was then retrieved from the obtained Hi-C matrices at 50 kbp for each condition (NC, C15m, and C4h) using hictkpy^76^ and numpy^85^ (Code: get_interaction_counts.py).

The non-metric multidimensional scaling (MDS) algorithm, isoMDS, from the MASS R package was used for dimensionality reduction with k = 2, to map the average 3D chromosomal distances onto a 2D plane. The output fitted coordinates were visualized using ggpubr and ggrepel.

Using hictk dump^76^, the .cool files of all samples were converted to bedpe files at 5mbp. The bedpe files were then read in R using DESeqDataSetFromMatrix and a differential contact analysis was performed using DESeq2 and standard parameters^64^. The differentially interacting interchromosomal domains were visualized using the Python package pyCirclize (https://github.com/moshi4/pyCirclize).

For each significant interchromosomal Hi-C interaction domain, we assigned an A/B compartment based on the greatest overlap with the domain’s genomic coordinates. A visualization was then created in Python to show the compartment-to-compartment interactions across different chromosomes. This plot was generated using the seaborn^86^ and matplotlib^70^ libraries (Supplemental Figures S8-S9). Gene coordinates were likewise overlapped with the significant interchromosomal Hi-C interaction domains, and the proportion of genes within each cluster that were present in these domains was plotted (Fig. 4D). Additionally, the respective domains on chromosome 9 were overlapped with the subcompartment domains and the share of each subcompartment is shown in Suppl. Fig. S7.

## Data and code access

All sequencing data generated in this study have been deposited in the European Nucleotide Archive (ENA; https://www.ebi.ac.uk/ena/) under the primary accession number PRJEB100712 and secondary accession number ERP182168. All codes used in the data analysis are available in our Git repository at https://github.com/paulsengroup/IMR90_mechano_confinement_2025/releases/tag/v1.0.1.

## Acknowledgments

This work was supported by grants from the Research Council of Norway (grants 351721, 343102 and 324137), the Norwegian Cancer Society (grant 223181), and Astri og Birger Torsteds legat.

Some of the analyses were performed on resources provided by Sigma2 – the National Infrastructure for High Performance Computing and Data Storage in Norway, with account number NN8041K.

Sequencing was performed by the Norwegian Sequencing Centre (https://www.sequencing.uio.no/), a national technology platform hosted by the University of Oslo and Oslo University Hospital and supported by Infrastructure programs of the Research Council of Norway and the Southeastern Regional Health Authorities.

We acknowledge the NorMIC Oslo imaging platform (Department of Biosciences, University of Oslo) and its technical staff We acknowledge the ENCODE Consortium and the ENCODE production laboratories of Bing Ren (University of California San Diego) and Michael Snyder (Stanford University).

Molecular graphics and analyses performed with UCSF ChimeraX, developed by the Resource for Biocomputing, Visualization, and Informatics at the University of California, San Francisco, with support from National Institutes of Health R01-GM129325 and the Office of Cyber Infrastructure and Computational Biology, National Institute of Allergy and Infectious Diseases.

## Authors’ contributions

CP and JP conceived and designed the study. SO and OH performed data analyses. SO performed Hi-C data analysis, and did downstream analyses including TEC clustering, 3D genome modeling, compartment and TAD analyses, MDS analyses and statistical analyses. OH did RNA-seq preprocessing. OH and AH did cell culture work. AH did imaging, immunostaining and cytokine assay analyses. CP and JP supervised the work. All authors read and approved the final manuscript.

