## Supplemental Figures for "The 3D nuclear position and compartmentalization of genes prime their response to mechano-confinement"

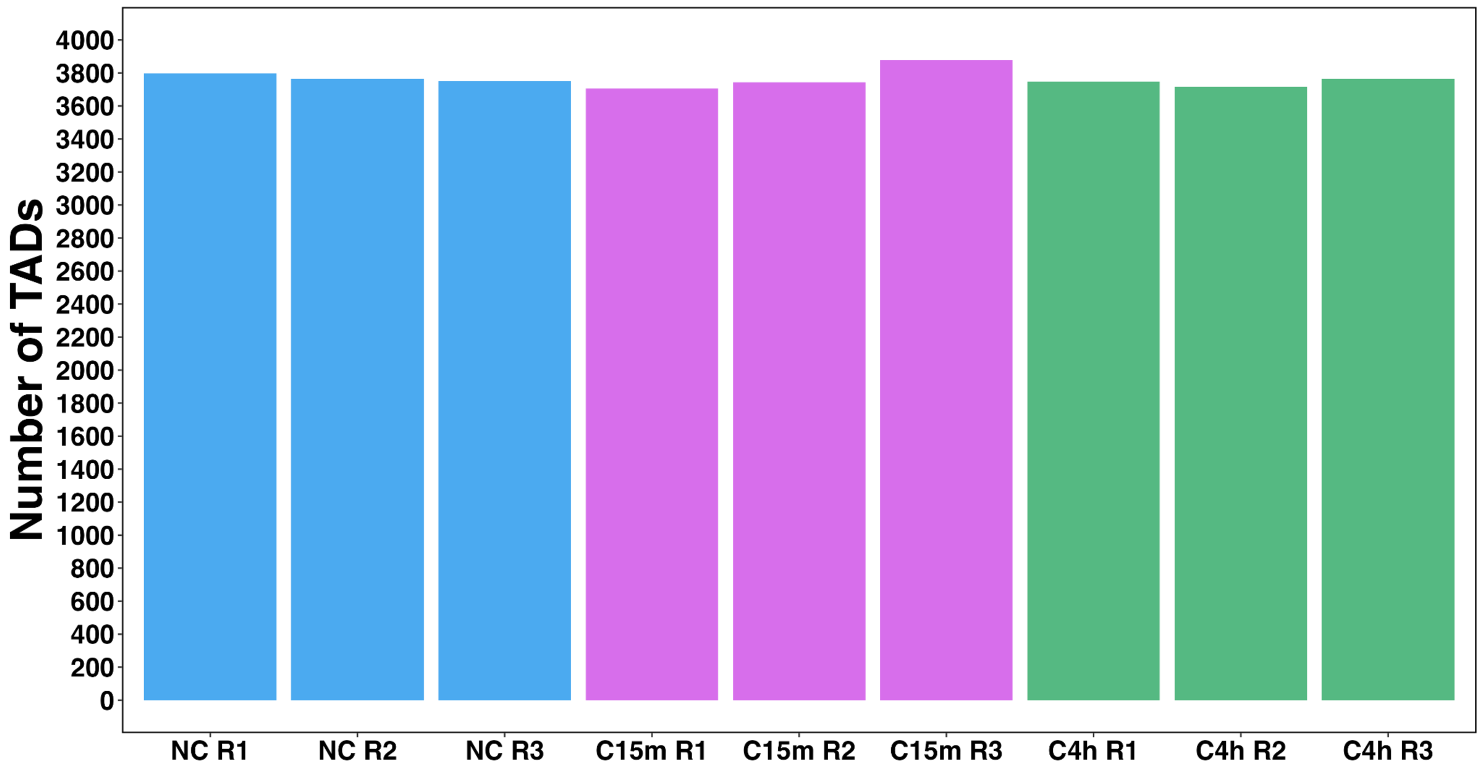


**Fig. S1:** The number of identified TADs in each replicate at 50kbp.

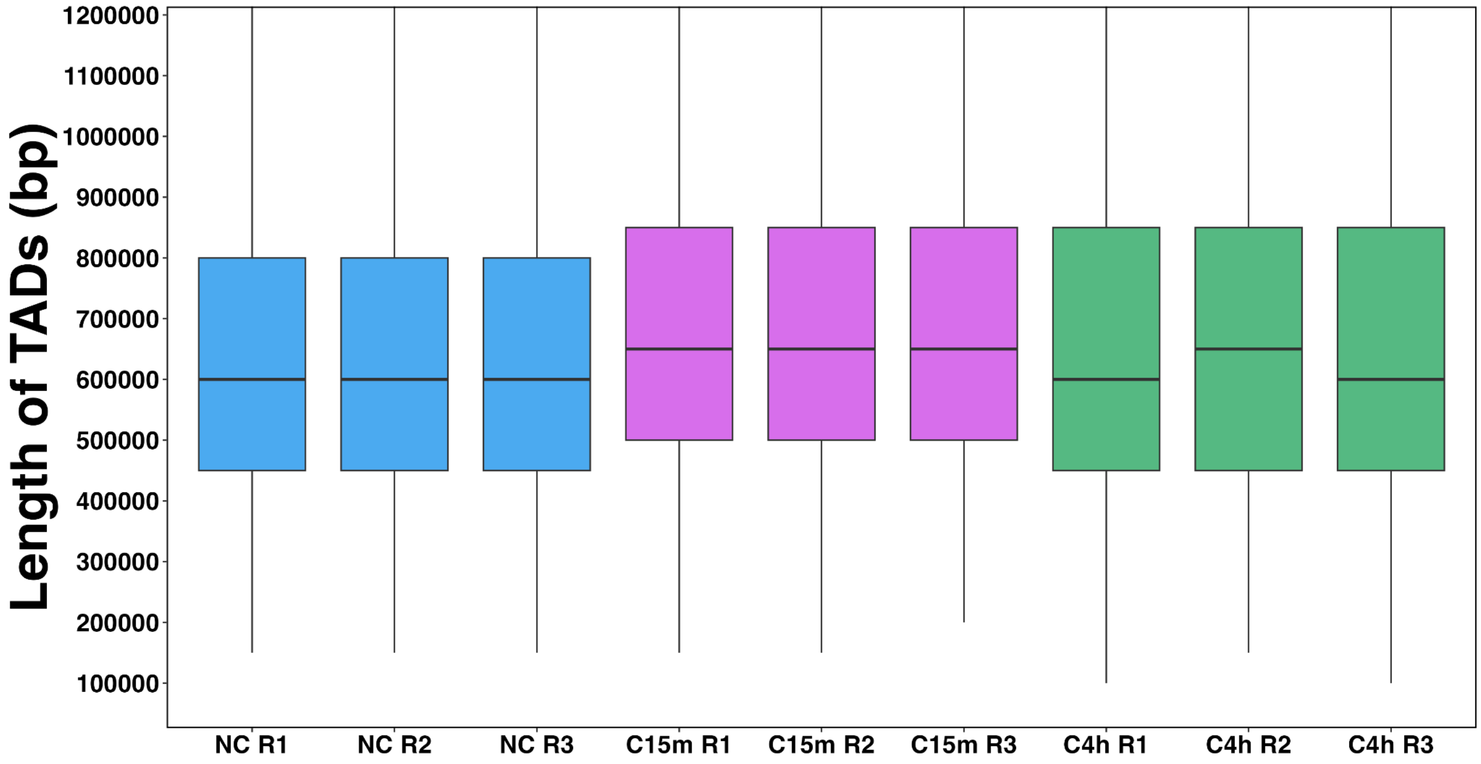


**Fig. S2:** Boxplot of the length of the identified TADs in each replicate at 50kbp.

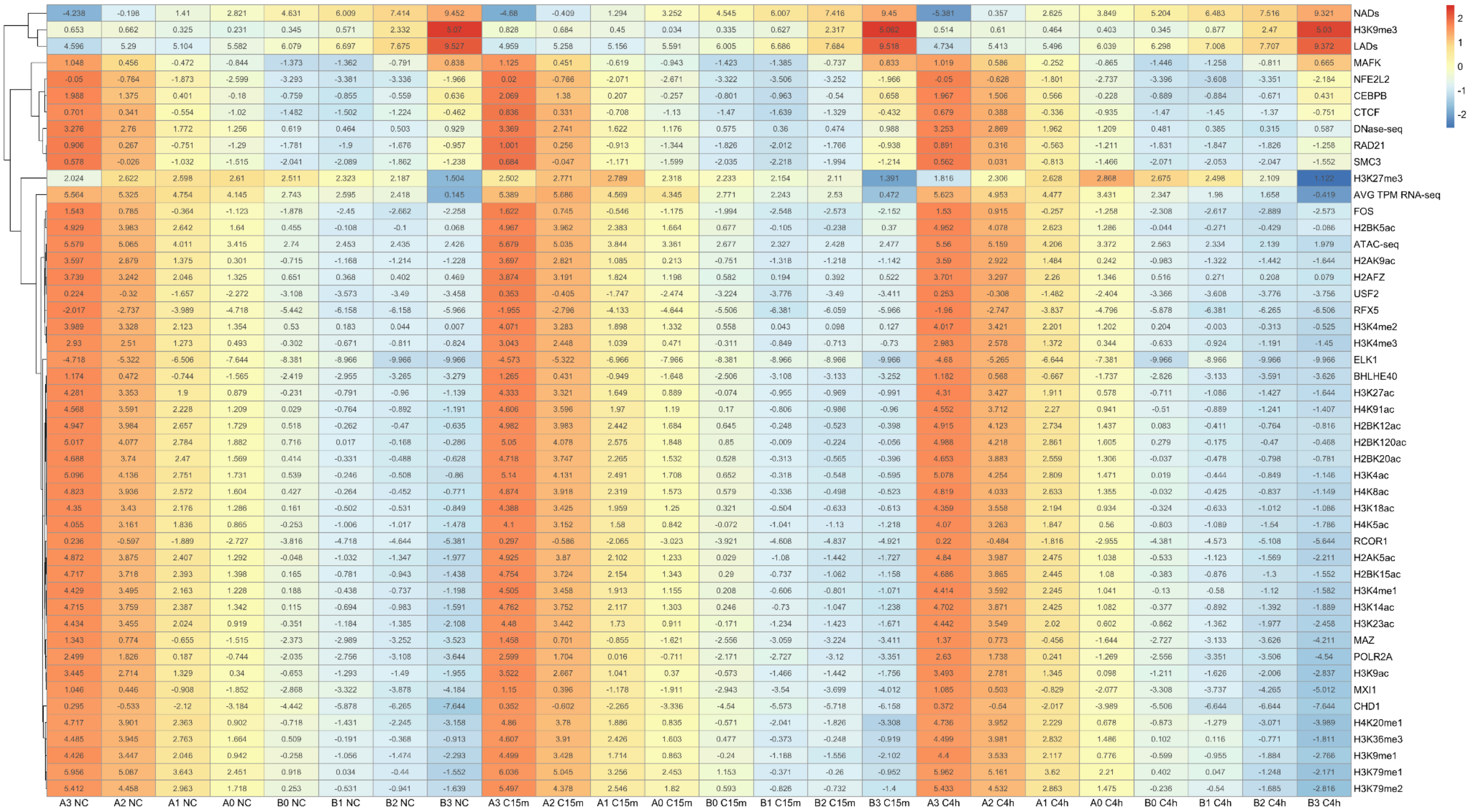


**Fig. S3:** Sub-compartment overlap with transcription factor and histone marker ChIP-seq data

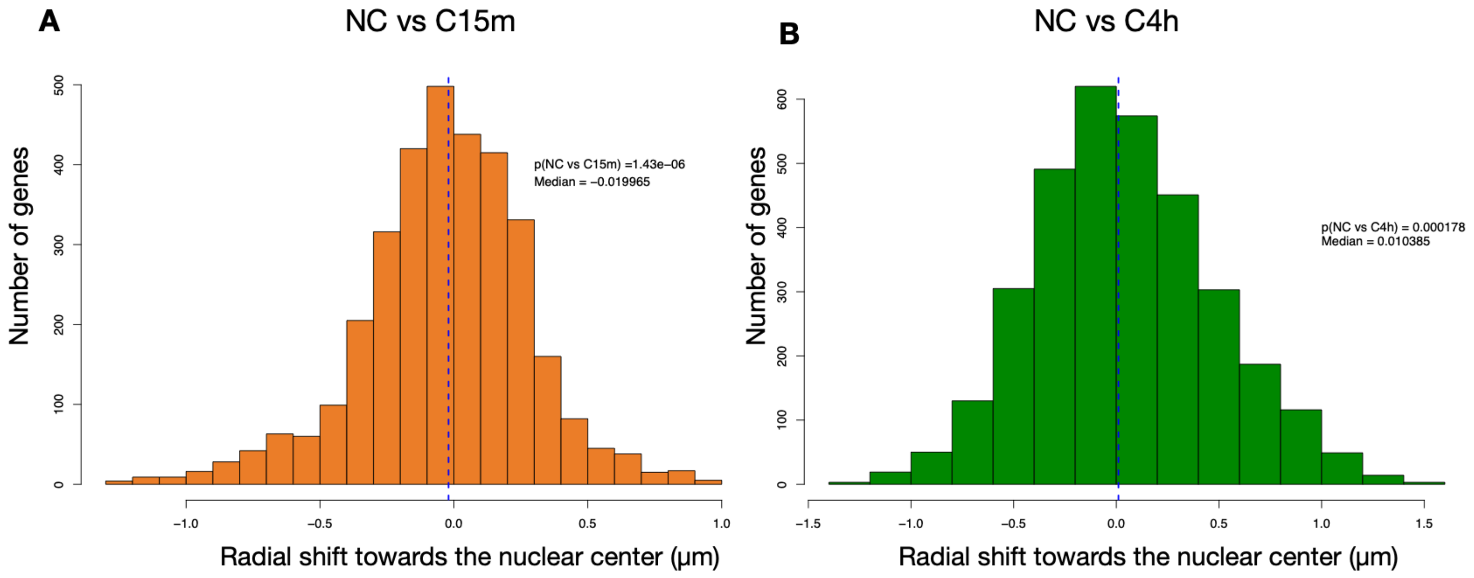


**Fig. S4:** Histogram showing the differences in median distances from the nuclear center for each TEC bead between NC vs. C15m (A) and NC vs. C4h (B) Chrom3D models. Negative and positive values indicate movement toward or away from the nuclear center, respectively. Pairwise Wilcoxon rank-sum tests were used to assess statistical significance of the differences in median distances between conditions.

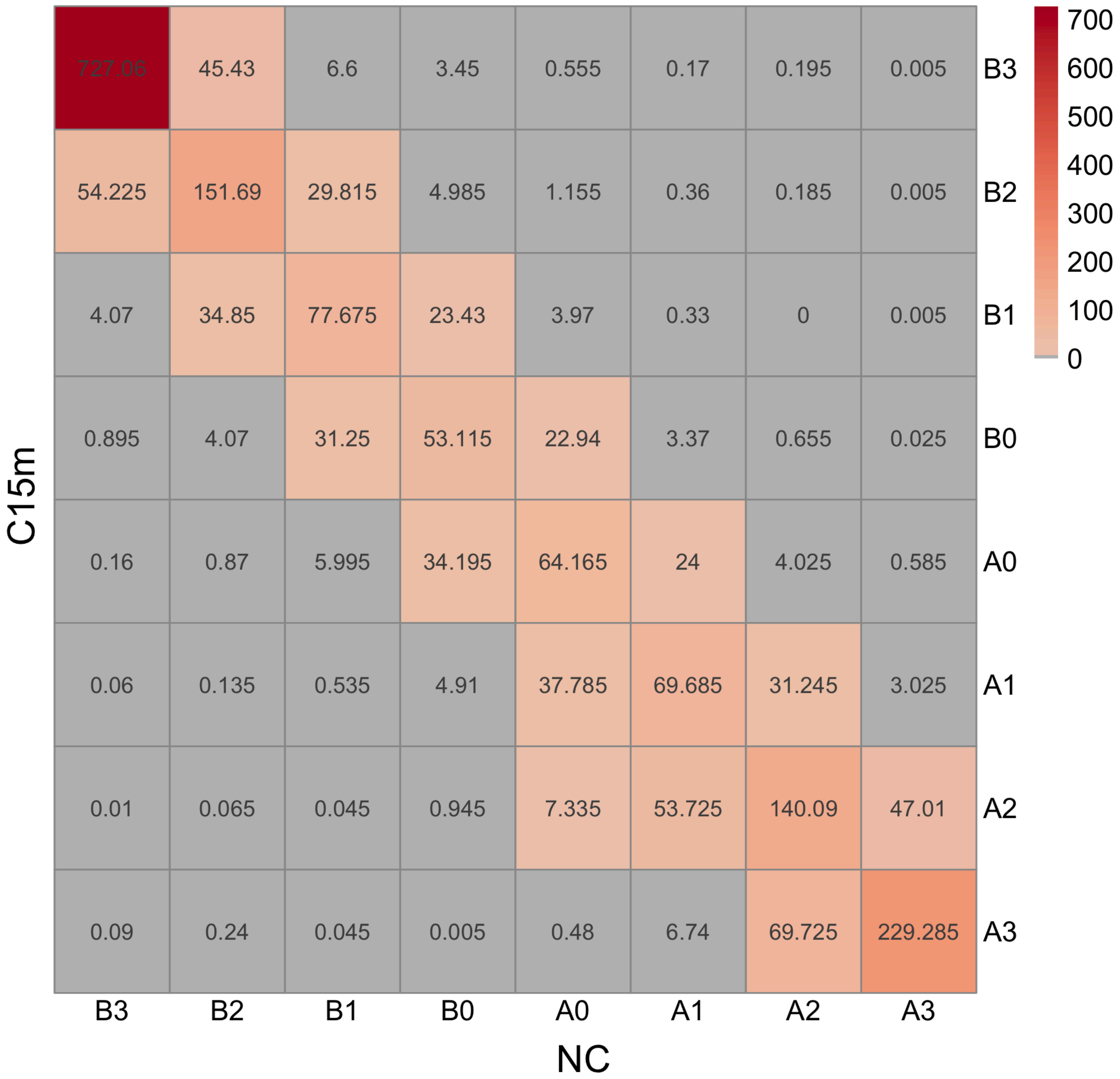


**Fig. S5:** Sub-compartment switches in the whole genome in Mbp NC vc. C15m

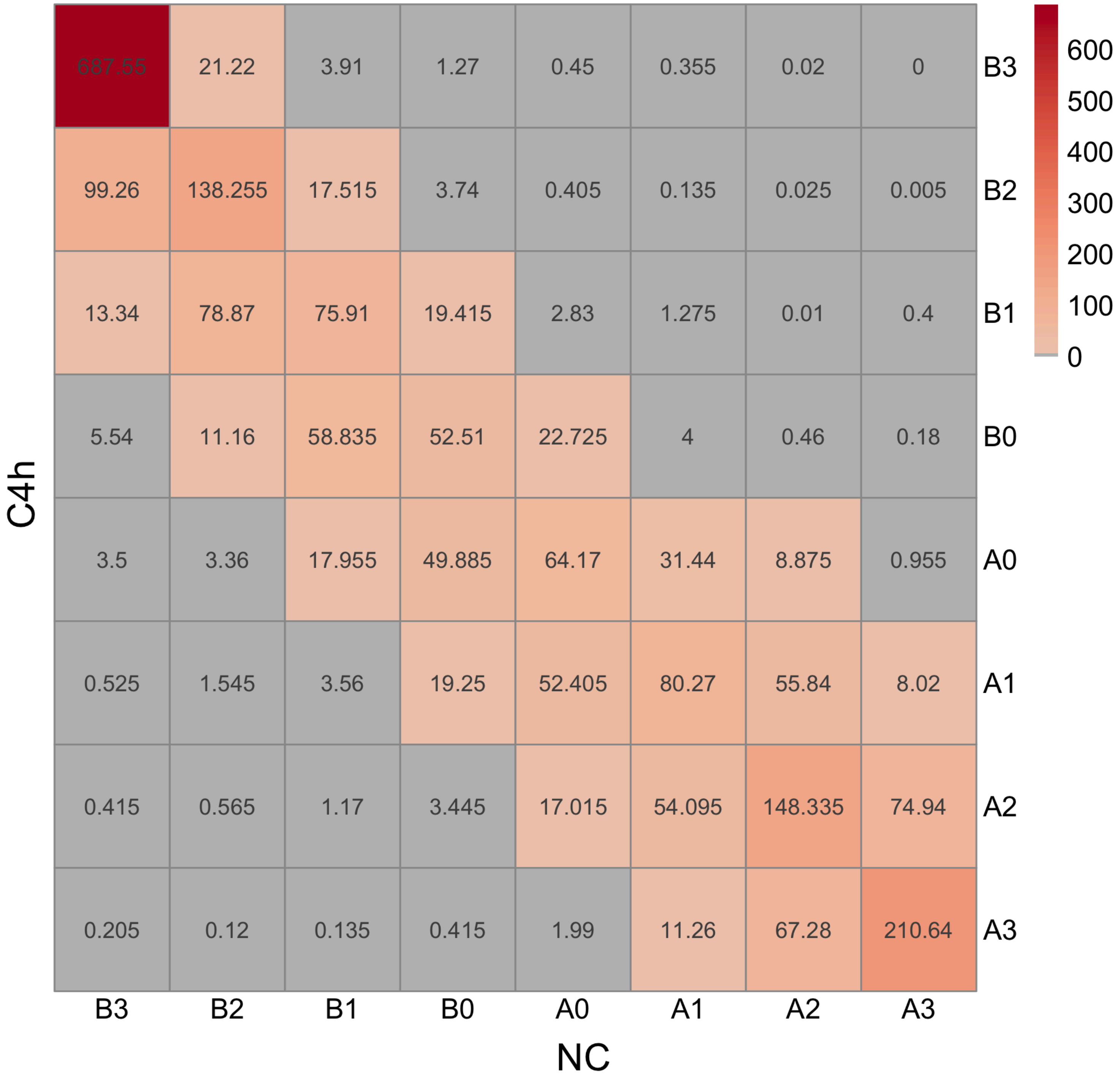


**Fig. S6:** Sub-compartment switches in the whole genome in Mbp NC vc. C4h

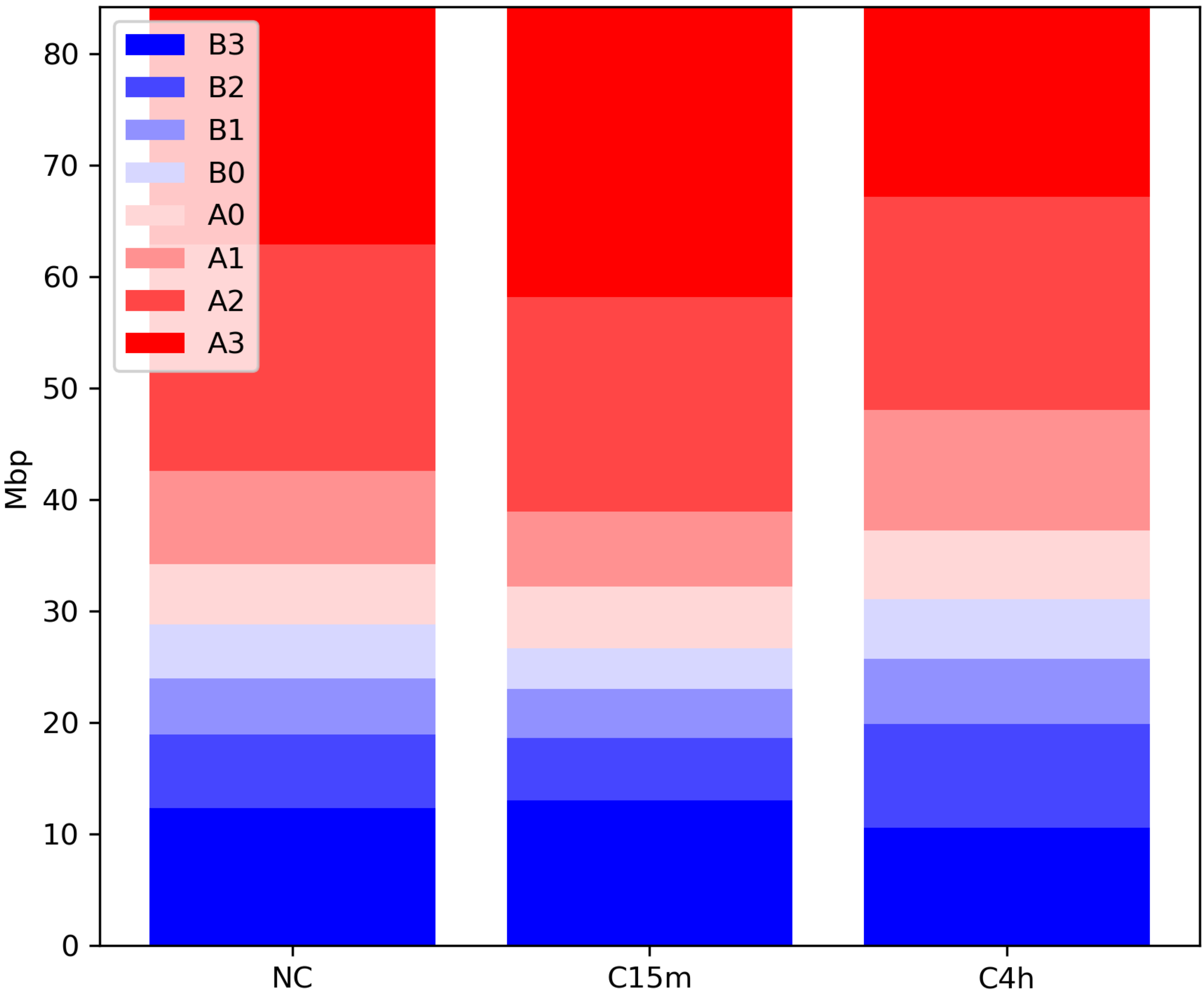


**Fig. S7:** Sub-compartment share in significant chromosome 9 inter-chromosomal interactions in NC vs. C15m

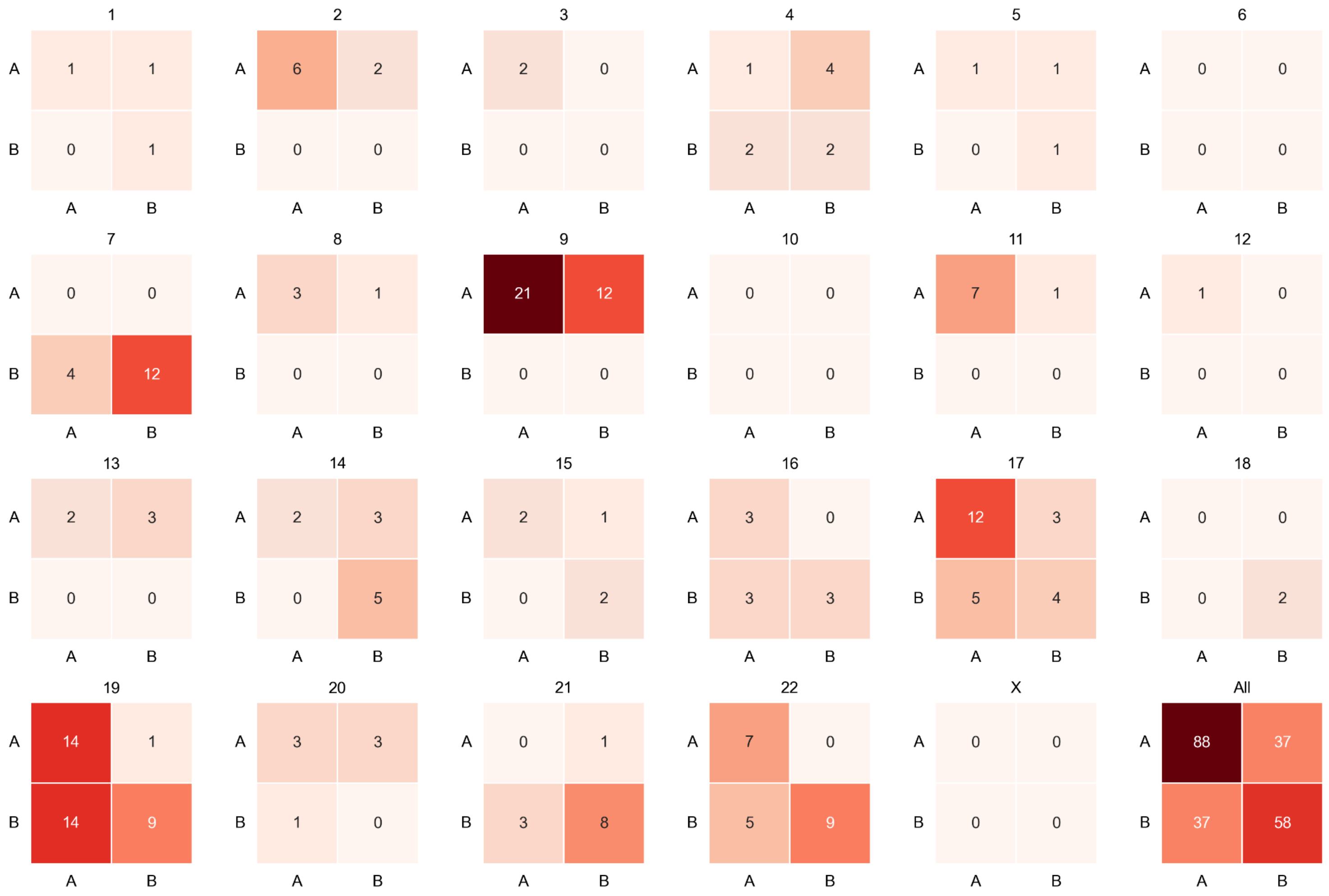


**Fig. S8:** Compartment status (A or B) for differentially interacting inter-chromosomal Hi-C regions in NC vs. C15m from each chromosome (indicated on top of each matrix), or jointly for all chromosomes (“All”).

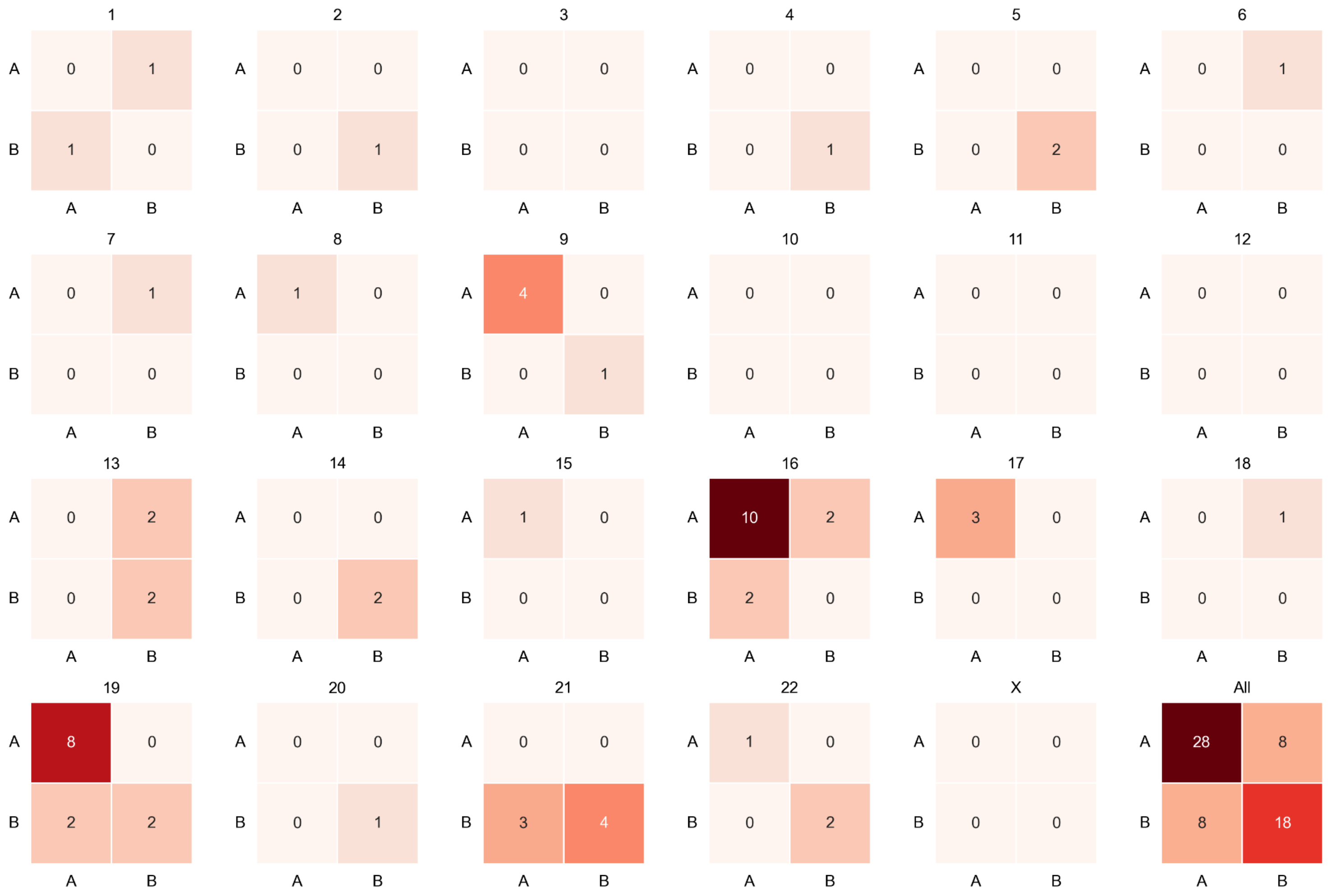


**Fig. S9:** Compartment status (A or B) for differentially interacting inter-chromosomal Hi-C regions in NC vs. C4h from each chromosome (indicated on top of each matrix), or jointly for all chromosomes (“All”).

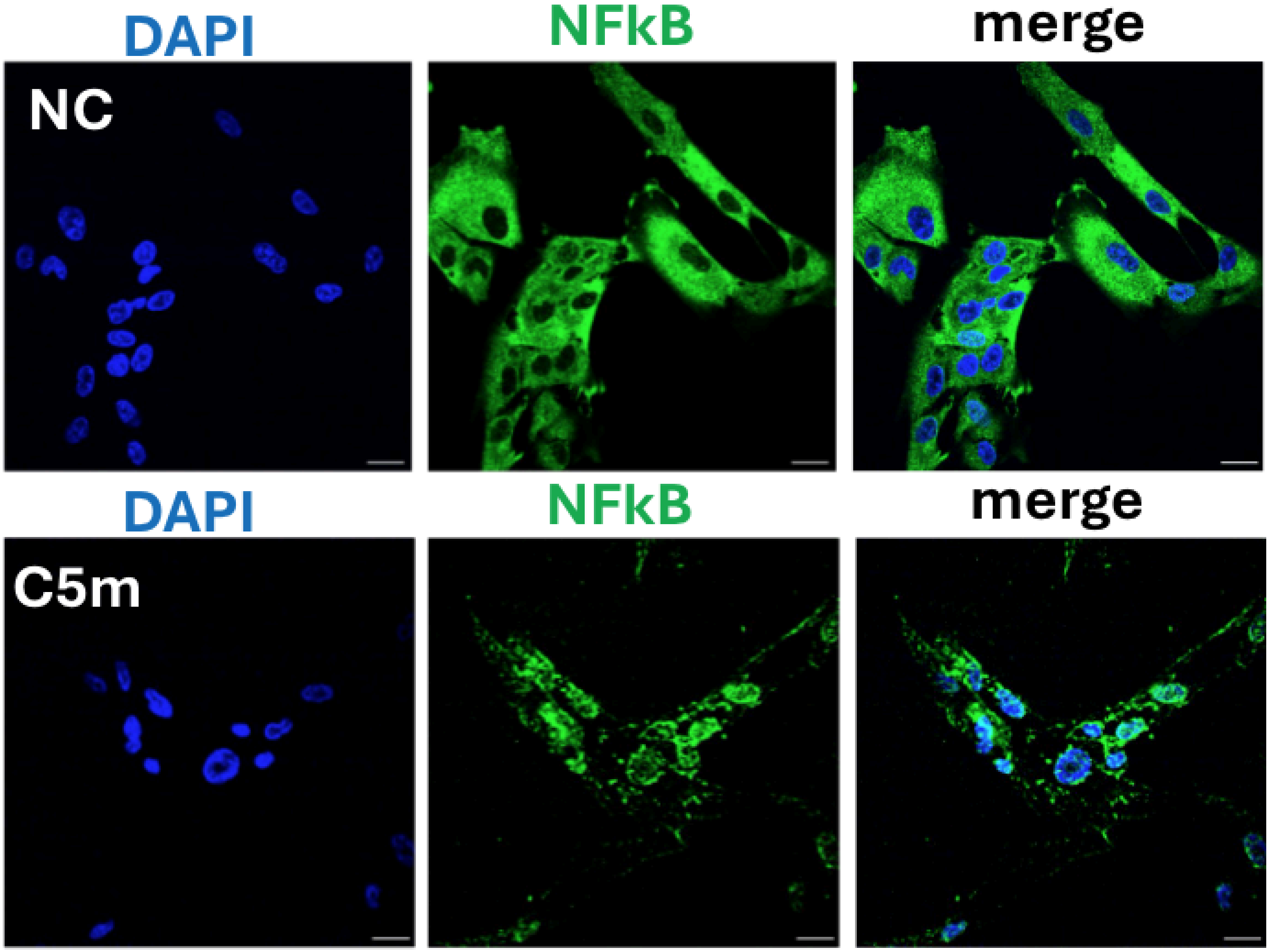


**Fig. S10:** Immunostaining of NFkB (green), and DAPI (blue) in NC (top) and cells confined for 5 minutes (C5m; bottom). Merge shown in the right panels.

**A**

| | 1 | 2 | 3 | 4 | 5 | 6 | 7 | 8 |
|---|---|---|---|---|---|---|---|---|
| A | NT-4 | IGFBP-3 | FGF-4 | Oncostatin M | MIP-1-delta | IL12-p40 | I-309 | Pos |
| B | Osteopontin | IGFBP-4 | FGF-6 | TPO | RANTES | IL-13 | IL-1alpha | Pos |
| C | Osteoprotegerin | IL-16 | FGF-7 | VEGF | SCF | IL-15 | IL-1beta | Pos |
| D | PARC | IP-10 | FGF-9 | PDGF-BB | SDF-1 | IFN-gamma | IL-2 | Pos |
| E | PIGF | LIF | Flt-3 Ligand | Leptin | TARC | MCP-1 | IL-3 | Neg |
| F | TGF- b 2 | LIGHT | Fractalkine | BDNF | TGF-beta 1 | MCP-2 | IL-4 | Neg |
| G | TGF- b 3 | MCP-4 | GCP-2 | BLC | TNF-alpha | MCP-3 | IL-5 | ENA-78 |
| H | TIMP-1 | MIF | GDNF | CK beta 8-1 | TNF-beta | M-CSF | IL-6 | G-CSF |
| I | TIMP-2 | MIP-3-alpha | HGF | Eotaxin | EGF | MDC | IL-7 | GM-CSF |
| J | Pos | NAP-2 | IGFBP-1 | Eotaxin-2 | IGF-1 | MIG | IL-8 | GRO |
| K | Pos | NT-3 | IGFBP-2 | Eotaxin-3 | Angiogenin | MIP-1 beta | IL-10 | GRO-alpha |

**B**

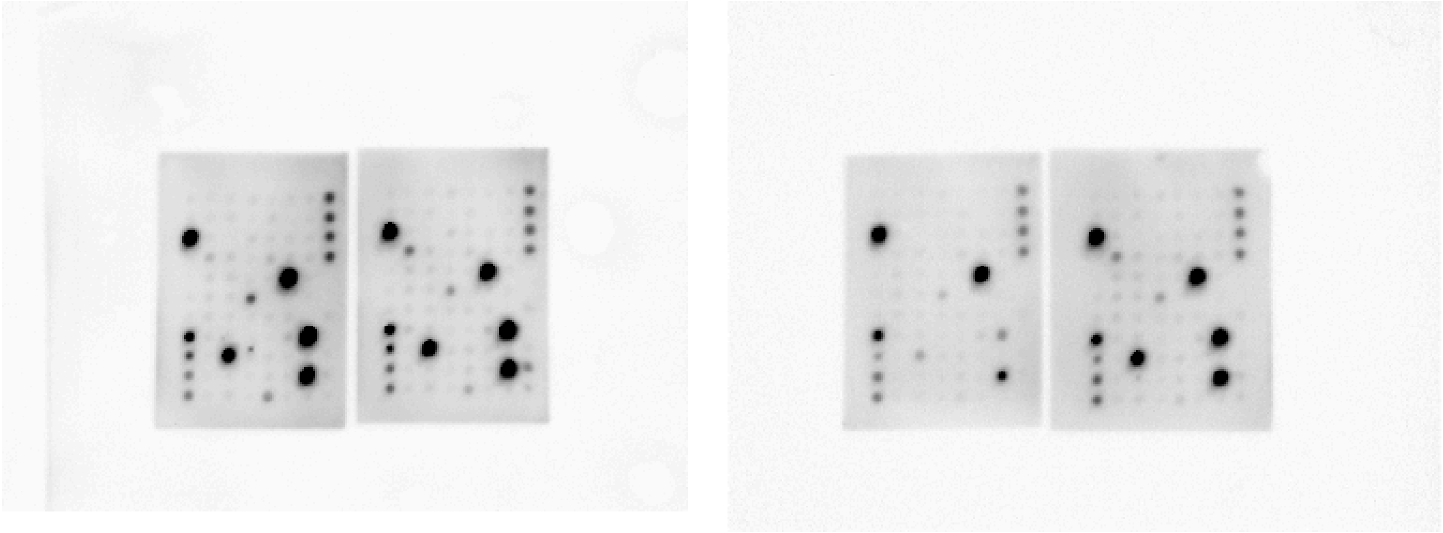

**Fig. S11: A:** Target map showing the assayed cytokines on the utilized C5 cytokine antibody array (RayBiotech, AAH-CYT-5) **B:** Images showing both replicates (left and right image), with each showing non-confined (NC) to the left, and confined at 4h (C4h) to the right.

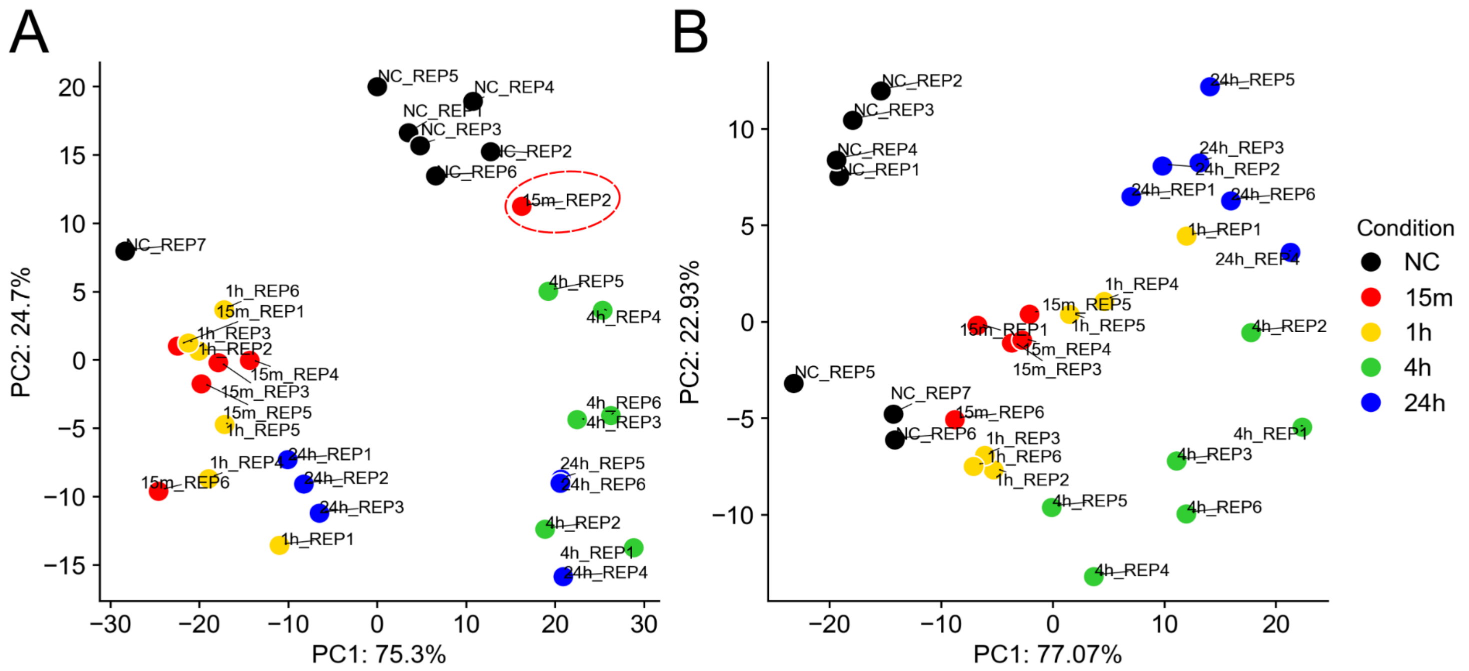


**Fig. S12:** Principal component analysis (PCA) of RNA-seq samples based on gene expression profiles. **A:** PCA plot of all sample replicates for each condition (shown in separate colors [see legend on the right]). The sample 15m_REP2 is highlighted by a red dotted circle. **B:** PCA plot after excluding sample 15m_REP2.
