## Supplemental Tables for "The 3D nuclear position and compartmentalization of genes prime their response to mechano-confinement"

**Table S1**: Sub-compartment switching composition by TEC NC vs. C15m

| TEC | 0 | | 1 | | 2 | | 3 | | 4 | | 5 | | 6 | | 7 | |
|---|---|---|---|---|---|---|---|---|---|---|---|---|---|---|---|---|
| **Total** | 6042 | | 5214 | | 2560 | | 2164 | | 356 | | 552 | | 213 | | 153 | |
| **Non-switch** | 2715 | 0.45 | 2390 | 0.46 | 1183 | 0.46 | 1032 | 0.48 | 172 | 0.48 | 244 | 0.44 | 89 | 0.42 | 73 | 0.48 |
| **Chromatin opening** | 1109 | 0.18 | 915 | 0.18 | 434 | 0.17 | 383 | 0.18 | 49 | 0.14 | 95 | 0.172 | 34 | 0.16 | 26 | 0.17 |
| **Chromatin closing** | 706 | 0.12 | 746 | 0.14 | 240 | 0.09 | 299 | 0.14 | 52 | 0.15 | 61 | 0.11 | 29 | 0.14 | 17 | 0.11 |
| **Facultative heterochromatin gain** | 215 | 0.036 | 255 | 0.05 | 37 | 0.01 | 66 | 0.030 | 23 | 0.06 | 9 | 0.0163 | 7 | 0.03 | 7 | 0.046 |
| **Facultative heterochromatin loss** | 252 | 0.04 | 254 | 0.05 | 65 | 0.03 | 77 | 0.035 | 13 | 0.037 | 18 | 0.033 | 6 | 0.028 | 7 | 0.046 |

**Table S2**: Sub-compartment switching composition by TEC NC vs. C4h

| TEC | 0 | | 1 | | 2 | | 3 | | 4 | | 5 | | 6 | | 7 | |
|---|---|---|---|---|---|---|---|---|---|---|---|---|---|---|---|---|
| **Total** | 6042 | | 5214 | | 2560 | | 2164 | | 356 | | 552 | | 213 | | 153 | |
| **Non-switch** | 2353 | 0.39 | 2067 | 0.4 | 1082 | 0.42 | 937 | 0.43 | 149 | 0.42 | 221 | 0.40 | 69 | 0.32 | 59 | 0.39 |
| **Chromatin opening** | 1352 | 0.22 | 1286 | 0.25 | 429 | 0.17 | 443 | 0.20 | 74 | 0.21 | 99 | 0.179 | 50 | 0.23 | 42 | 0.27 |
| **Chromatin closing** | 825 | 0.14 | 698 | 0.13 | 346 | 0.14 | 334 | 0.15 | 50 | 0.14 | 80 | 0.14 | 33 | 0.15 | 15 | 0.098 |
| **Facultative heterochromatin gain** | 337 | 0.06 | 317 | 0.06 | 80 | 0.031 | 89 | 0.041 | 23 | 0.065 | 24 | 0.043 | 9 | 0.04 | 11 | 0.072 |
| **Facultative heterochromatin loss** | 315 | 0.05 | 362 | 0.07 | 70 | 0.027 | 105 | 0.049 | 21 | 0.059 | 19 | 0.034 | 14 | 0.066 | 11 | 0.072 |

**Table S3**: Mean log2FC per cluster and condition, rounded to 3 decimal places

| TEC | C15m | C1h | C4h | C24h |
|---|---|---|---|---|
| 0 | 0.008 | 0.309 | 0.038 | 0.086 |
| 1 | 0.396 | -0.398 | -0.172 | -0.25 |
| 2 | -0.851 | 1.184 | 0.443 | 0.63 |
| 3 | -0.206 | 0.393 | -0.118 | 0.033 |
| 4 | 2.14 | -2.16 | -0.858 | -1.73 |
| 5 | -1.28 | 4.008 | 0.247 | 1.038 |
| 6 | 0.405 | -0.434 | 1.504 | 1.423 |
| 7 | -1.574 | 1.246 | 3.291 | 3.54 |

**Table S4:** Hi-C interaction statistics by replicate and condition

| | NC_rep1 | NC_rep2 | NC_rep3 | C15m_rep1 | C15m_rep2 | C15m_rep3 | C4h_rep1 | C4h_rep2 | C4h_rep3 |
|---|---|---|---|---|---|---|---|---|---|
| valid_interaction | 481153485 | 566639409 | 610851332 | 554462543 | 599472699 | 697116600 | 555197933 | 702914209 | 547940609 |
| valid_interaction_rmdup | 402519041 | 477289162 | 497843748 | 386142107 | 365213766 | 454286864 | 459671694 | 581461162 | 456636315 |
| trans_interaction | 61881584 | 64460192 | 53305729 | 54082262 | 62963926 | 73608358 | 80671856 | 118613919 | 89725395 |
| cis_interaction | 340637457 | 412828970 | 444538019 | 332059845 | 302249840 | 380678506 | 378999838 | 462847243 | 366910920 |
| cis_shortRange | 131738738 | 156334274 | 180647614 | 131871725 | 107109512 | 172266877 | 141944933 | 145993135 | 143407393 |
| cis_longRange | 208898719 | 256494696 | 263890405 | 200188120 | 195140328 | 208411629 | 237054905 | 316854108 | 223503527 |
| Valid_interaction_pairs_FF | 112015871 | 132063886 | 139957461 | 123351464 | 137760899 | 145493640 | 126702723 | 165113372 | 123845440 |
| Valid_interaction_pairs_RR | 112092525 | 132109281 | 140096374 | 123443846 | 137882037 | 145596517 | 126726396 | 165245739 | 123888339 |
| Valid_interaction_pairs_RF | 109845362 | 129673457 | 136543804 | 121680370 | 136430673 | 140980091 | 123557171 | 163363331 | 120558892 |
| Valid_interaction_pairs_FR | 147199727 | 172792785 | 194253693 | 185986863 | 187399090 | 265046352 | 178211643 | 209191767 | 179647938 |
| Dangling_end_pairs | 3877003 | 2810313 | 5121418 | 1874453 | 2053604 | 2812663 | 4105637 | 4917308 | 7797807 |
| Religation_pairs | 7764531 | 6924555 | 11143466 | 5143712 | 5070752 | 7621754 | 8671069 | 9854691 | 13816127 |
| Self_Cycle_pairs | 4219 | 5860 | 5888 | 10222 | 14827 | 17347 | 6345 | 4382 | 4812 |
| Single-end_pairs | 0 | 0 | 0 | 0 | 0 | 0 | 0 | 0 | 0 |
| Filtered_pairs | 0 | 0 | 0 | 0 | 0 | 0 | 0 | 0 | 0 |
| Dumped_pairs | 1217 | 1177 | 1507 | 706 | 710 | 1034 | 1435 | 993 | 1527 |

**Table S5**: Sub-compartment share by condition and replicate in percentage

| sub_com | NC_rep1 | NC_rep2 | NC_rep3 | C15min_rep1 | C15min_rep2 | C15min_rep3 | C4h_rep1 | C4h_rep2 | C4h_rep3 |
|---|---|---|---|---|---|---|---|---|---|
| A3 | 11.29 | 9.72 | 10.69 | 10.79 | 10.98 | 11.8 | 10.08 | 11.5 | 10.94 |
| A2 | 11.77 | 10.81 | 11.59 | 11.42 | 11.42 | 10.82 | 12.47 | 12.59 | 13.29 |
| A1 | 9.45 | 8.76 | 9.06 | 8.67 | 8.35 | 7.49 | 10.77 | 10.62 | 11.42 |
| A0 | 8.13 | 7.96 | 8.77 | 7.47 | 8.57 | 7.08 | 9.01 | 9.33 | 9.51 |
| B0 | 7.46 | 8.45 | 8.03 | 7.22 | 7.33 | 6.22 | 8.34 | 7.47 | 7.65 |
| B1 | 8.58 | 8.96 | 8.59 | 7.41 | 7.58 | 7.93 | 9.21 | 8.32 | 7.89 |
| B2 | 10.32 | 10.42 | 10.7 | 9.6 | 9.49 | 12 | 10.28 | 9.52 | 10.14 |
| B3 | 26.95 | 28.91 | 26.6 | 26.84 | 25.69 | 26.68 | 23.92 | 24.84 | 23.22 |
| NA | 6.04 | 6.01 | 5.99 | 10.58 | 10.61 | 9.96 | 5.93 | 5.81 | 5.94 |

**Table S6**: Sub-compartment share by condition and replicate in Mbp

| sub_com | NC_rep1 | NC_rep2 | NC_rep3 | C15min_rep1 | C15min_rep2 | C15min_rep3 | C4h_rep1 | C4h_rep2 | C4h_rep3 |
|---|---|---|---|---|---|---|---|---|---|
| A3 | 342.265 | 294.72 | 323.905 | 327.175 | 332.725 | 357.735 | 305.58 | 348.435 | 331.59 |
| A2 | 356.675 | 327.75 | 351.19 | 346.235 | 346.05 | 327.92 | 377.905 | 381.665 | 402.815 |
| A1 | 286.575 | 265.43 | 274.555 | 262.68 | 253.085 | 227.13 | 326.445 | 322.035 | 346.025 |
| A0 | 246.41 | 241.315 | 265.775 | 226.31 | 259.665 | 214.61 | 273.045 | 282.78 | 288.23 |
| B0 | 226.015 | 256.025 | 243.275 | 218.955 | 222.035 | 188.66 | 252.815 | 226.53 | 231.775 |
| B1 | 260.15 | 271.515 | 260.24 | 224.615 | 229.82 | 240.505 | 279.095 | 252.215 | 239.275 |
| B2 | 312.96 | 315.83 | 324.3 | 290.985 | 287.58 | 363.805 | 311.655 | 288.495 | 307.48 |
| B3 | 816.935 | 876.365 | 806.31 | 813.6 | 778.64 | 808.84 | 724.905 | 752.805 | 703.805 |
| NA | 183.11 | 182.145 | 181.545 | 320.54 | 321.495 | 301.89 | 179.65 | 176.135 | 180.1 |

**Table S7:** Median distance from the nucleus center and the standard deviation in Chrom3D models per condition and subcompartment

| subcompartment | median distance ± std - NC | median distance ± std - C15m | median distance ± std - C4h |
|---|---|---|---|
| A3 | 3.35 ± 0.33 | 3.24 ± 0.43 | 3.43 ± 0.37 |
| A2 | 3.17 ± 0.33 | 3.04 ± 0.44 | 3.35 ± 0.33 |
| A1 | 3.30 ± 0.32 | 3.33 ± 0.46 | 3.43 ± 0.33 |
| A0 | 3.42 ± 0.32 | 3.60 ± 0.42 | 3.49 ± 0.36 |
| B0 | 3.60 ± 0.32 | 3.45 ± 0.40 | 3.62 ± 0.34 |
| B1 | 3.65 ± 0.32 | 3.64 ± 0.42 | 3.66 ± 0.32 |
| B2 | 3.64 ± 0.30 | 3.66 ± 0.40 | 3.73 ± 0.35 |
| B3 | 3.75 ± 0.29 | 3.62 ± 0.40 | 3.70 ± 0.31 |

**Table S8**: pairwise Wilcox results of the median subcompartment distance from the nucleus center per sub-compartment within NC Chrom3D simulations

| | A3 | A2 | A1 | A0 | B0 | B1 | B2 |
|---|---|---|---|---|---|---|---|
| A3 | | | | | | | |
| A2 | 4.54e-03 | | | | | | |
| A1 | 1.00e+00 | 1.61e-02 | | | | | |
| A0 | 6.02e-01 | 1.24e-05 | 2.73e-01 | | | | |
| B0 | 7.17e-08 | 8.23e-16 | 2.19e-09 | 9.71e-05 | | | |
| B1 | 8.47e-09 | 1.10e-16 | 1.87e-10 | 8.53e-06 | 1.00e+00 | | |
| B2 | 8.86e-10 | 4.76e-18 | 2.23e-11 | 2.26e-06 | 1.00e+00 | 1.00e+00 | |
| B3 | 7.12e-15 | 6.30e-23 | 6.54e-17 | 2.73e-11 | 5.00e-03 | 7.23e-02 | 8.82e-02 |

**Table S9**: pairwise Wilcox results of the median cluster distance from the nucleus center per sub-compartment within C15m Chrom3D simulations

| | A3 | A2 | A1 | A0 | B0 | B1 | B2 |
|---|---|---|---|---|---|---|---|
| A3 | | | | | | | |
| A2 | 1.82e-04 | | | | | | |
| A1 | 4.93e-01 | 5.58e-07 | | | | | |
| A0 | 1.61e-10 | 4.57e-16 | 4.08e-07 | | | | |
| B0 | 7.61e-06 | 2.76e-12 | 7.70e-03 | 2.39e-02 | | | |
| B1 | 7.75e-12 | 5.83e-17 | 2.55e-08 | 1.00e+00 | 2.81e-03 | | |
| B2 | 3.38e-13 | 4.22e-18 | 4.81e-10 | 4.93e-01 | 6.71e-05 | 1.00e+00 | |
| B3 | 3.95e-11 | 1.27e-16 | 1.45e-07 | 1.00e+00 | 1.66e-02 | 1.00e+00 | 4.93e-01 |

**Table S10**: pairwise Wilcox results of the median cluster distance from the nucleus center per sub-compartment within C4h Chrom3D simulations

| | A3 | A2 | A1 | A0 | B0 | B1 | B2 |
|---|---|---|---|---|---|---|---|
| A3 | | | | | | | |
| A2 | 2.56e-01 | | | | | | |
| A1 | 1.00e+00 | 4.97e-01 | | | | | |
| A0 | 1.00e+00 | 5.56e-03 | 5.87e-01 | | | | |
| B0 | 9.43e-05 | 7.63e-09 | 1.89e-05 | 7.07e-03 | | | |
| B1 | 1.00e-07 | 9.67e-12 | 3.78e-08 | 2.72e-05 | 7.37e-01 | | |
| B2 | 1.06e-08 | 6.01e-13 | 1.98e-09 | 3.85e-06 | 2.56e-01 | 1.00e+00 | |
| B3 | 1.20e-08 | 5.46e-13 | 1.76e-09 | 3.01e-06 | 2.08e-01 | 1.00e+00 | 1.00e+00 |

**Table S11**: Wilcoxon rank sum test results of the median subcompartment distance from the nucleus center in each Chrom3D simulation model per condition and subcompartment pair.

| Condition 1 | Condition 2 | Subcomp. | P-value |
|---|---|---|---|
| NC | C15m | A3 | 0.509 |
| NC | C15m | A2 | 0.185 |
| NC | C15m | A1 | 0.172 |
| NC | C15m | A0 | 2.20 × 10^-07 |
| NC | C15m | B0 | 0.040 |
| NC | C15m | B1 | 0.329 |
| NC | C15m | B2 | 0.122 |
| NC | C15m | B3 | 0.008 |
| NC | C4h | A3 | 0.002 |
| NC | C4h | A2 | 2.28 × 10^-06 |
| NC | C4h | A1 | 5.89 × 10^-04 |
| NC | C4h | A0 | 0.003 |
| NC | C4h | B0 | 0.193 |
| NC | C4h | B1 | 0.069 |
| NC | C4h | B2 | 0.026 |
| NC | C4h | B3 | 0.619 |
| C15m | C4h | A3 | 3.82 × 10^-05 |
| C15m | C4h | A2 | 5.04 × 10^-09 |
| C15m | C4h | A1 | 0.043 |
| C15m | C4h | A0 | 0.003 |
| C15m | C4h | B0 | 6.22 × 10^-04 |
| C15m | C4h | B1 | 0.392 |
| C15m | C4h | B2 | 0.509 |
| C15m | C4h | B3 | 0.011 |

**Table S12:** Median distance from the nucleus center and the standard deviation in Chrom3D models per condition and TEC

| TEC | median distance ± std - NC | median distance ± std - C15m | median distance ± std - C4h |
|---|---|---|---|
| 0 | 3.66 ± 0.32 | 3.62 ± 0.42 | 3.72 ± 0.36 |
| 1 | 3.71 ± 0.32 | 3.60 ± 0.44 | 3.70 ± 0.35 |
| 2 | 3.06 ± 0.31 | 2.96 ± 0.43 | 3.06 ± 0.33 |
| 3 | 3.37 ± 0.31 | 3.31 ± 0.41 | 3.44 ± 0.32 |
| 4 | 3.62 ± 0.32 | 3.54 ± 0.42 | 3.66 ± 0.34 |
| 5 | 3.16 ± 0.33 | 3.12 ± 0.43 | 3.21 ± 0.35 |
| 6 | 3.30 ± 0.31 | 3.25 ± 0.40 | 3.40 ± 0.34 |
| 7 | 3.48 ± 0.35 | 3.37 ± 0.45 | 3.48 ± 0.42 |

**Table S13**: pairwise Wilcox results of the median cluster distance from the nucleus center per cluster within NC Chrom3D simulations

| | 0 | 1 | 2 | 3 | 4 | 5 | 6 |
|---|---|---|---|---|---|---|---|
| 0 | | | | | | | |
| 1 | 3.82e-01 | | | | | | |
| 2 | 6.55e-23 | 4.03e-24 | | | | | |
| 3 | 4.61e-09 | 1.01e-10 | 1.06e-10 | | | | |
| 4 | 3.25e-01 | 1.50e-01 | 2.00e-20 | 6.89e-06 | | | |
| 5 | 1.72e-18 | 6.88e-20 | 1.47e-01 | 1.42e-05 | 1.74e-15 | | |
| 6 | 8.62e-13 | 1.13e-14 | 7.35e-07 | 2.04e-01 | 1.43e-09 | 1.10e-02 | |
| 7 | 7.06e-04 | 3.72e-05 | 2.81e-14 | 1.50e-01 | 5.64e-02 | 4.61e-09 | 1.16e-03 |

**Table S14**: pairwise Wilcox results of the median cluster distance from the nucleus center per cluster within C15m Chrom3D simulations

| | 0 | 1 | 2 | 3 | 4 | 5 | 6 |
|---|---|---|---|---|---|---|---|
| 0 | | | | | | | |
| 1 | 7.42e-01 | | | | | | |
| 2 | 8.75e-20 | 2.87e-20 | | | | | |
| 3 | 2.68e-08 | 3.38e-09 | 3.16e-10 | | | | |
| 4 | 3.91e-01 | 3.01e-01 | 8.17e-18 | 3.31e-06 | | | |
| 5 | 3.03e-15 | 6.40e-16 | 9.96e-03 | 2.83e-05 | 3.24e-13 | | |
| 6 | 1.75e-11 | 2.11e-12 | 2.54e-07 | 3.04e-01 | 5.09e-09 | 9.52e-03 | |
| 7 | 3.63e-05 | 9.29e-06 | 1.75e-11 | 4.46e-01 | 3.11e-03 | 1.63e-06 | 2.90e-02 |

**Table S15**: pairwise Wilcox results of the median cluster distance from the nucleus center per cluster within C4h Chrom3D simulations

| | 0 | 1 | 2 | 3 | 4 | 5 | 6 |
|---|---|---|---|---|---|---|---|
| 0 | | | | | | | |
| 1 | 9.39e-01 | | | | | | |
| 2 | 3.93e-24 | 4.90e-24 | | | | | |
| 3 | 9.71e-10 | 7.34e-10 | 1.18e-13 | | | | |
| 4 | 4.01e-01 | 4.01e-01 | 3.79e-22 | 1.50e-06 | | | |
| 5 | 8.70e-19 | 9.79e-19 | 3.82e-03 | 1.86e-07 | 1.18e-16 | | |
| 6 | 1.46e-12 | 8.46e-13 | 3.12e-10 | 2.81e-01 | 1.32e-09 | 5.85e-04 | |
| 7 | 3.19e-05 | 1.85e-05 | 8.37e-16 | 4.01e-01 | 3.82e-03 | 4.91e-10 | 8.54e-03 |

**Table S16**: Wilcoxon rank sum test results of the median cluster distance from the nucleus center in each Chrom3D simulation model per condition and subcompartment pair.

| Condition 1 | Condition 2 | TEC | P-value |
|---|---|---|---|
| NC | C15m | 0 | 0.537 |
| NC | C15m | 1 | 0.273 |
| NC | C15m | 2 | 0.191 |
| NC | C15m | 3 | 0.517 |
| NC | C15m | 4 | 0.460 |
| NC | C15m | 5 | 0.776 |
| NC | C15m | 6 | 0.614 |
| NC | C15m | 7 | 0.227 |
| NC | C4h | 0 | 0.073 |
| NC | C4h | 1 | 0.392 |
| NC | C4h | 2 | 0.755 |
| NC | C4h | 3 | 0.031 |
| NC | C4h | 4 | 0.088 |
| NC | C4h | 5 | 0.151 |
| NC | C4h | 6 | 0.024 |
| NC | C4h | 7 | 0.177 |
| C15m | C4h | 0 | 0.014 |
| C15m | C4h | 1 | 0.035 |
| C15m | C4h | 2 | 0.095 |
| C15m | C4h | 3 | 0.003 |
| C15m | C4h | 4 | 0.011 |
| C15m | C4h | 5 | 0.063 |
| C15m | C4h | 6 | 0.007 |
| C15m | C4h | 7 | 0.007 |

**Table S17:** Fisher's exact test results of chromatin opening vs closing in upregulated clusters (TEC 2, 5, 6, 7) vs. downregulated TECs (TEC 1, 3, 4)

P-value = 3.805e-05; Odds ratio = 0.7234678

| | Chromatin closing in C15m | Chromatin opening in C15m |
|---|---|---|
| Upregulated TECs | 347 | 589 |
| Downregulated TECs | 1097 | 1347 |

P-value = 0.0008173; Odds ratio 1.27386

| | Chromatin closing in C4h | Chromatin opening in C4h |
|---|---|---|
| Upregulated TECs | 474 | 620 |
| Downregulated TECs | 1082 | 1803 |

**Table S18:** Cellular localization of p-NF-kB in IMR90 cells (n.o. cells).*

| | Condition: | NC | C5m |
|---|---|---|---|
| Localization: | Cytoplasm | 107 | 2 |
| | Nucleus | 3 | 139 |

* Fisher’s exact test of [NC vs. C5m] vs. [Cytoplasm vs. Nucleus] gives P<2.2e-16

**Table S19:** Cellular localization of NF-kB in IMR90 cells (n.o. cells).*

| | Condition: | NC | C5m |
|---|---|---|---|
| Localization: | Cytoplasm | 123 | 3 |
| | Nucleus | 1 | 104 |

* Fisher’s exact test of [NC vs. C5m] vs. [Cytoplasm vs. Nucleus] gives P<2.2e-16

**Table S20:** Intensity values from two replicates using the cytokine antibody array (RayBiotech, AAH-CYT-5).

| Factor | Rep1 (intensity) | Rep2 (intensity) | Avg. (intensity) | S.D. |
|---|---|---|---|---|
| TIMP1 | 1.75416733 | 1.25902193 | 1.50659463 | 0.35012067 |
| TIMP2 | 2.00292586 | 1.55761751 | 1.78027169 | 0.31488056 |
| OPG | 1.72867408 | 1.06321263 | 1.39594335 | 0.4705523 |
| CXCL10 | 4.7924873 | 2.86982663 | 3.83115697 | 1.35952639 |
| HGF | 1.78113965 | 6.57749225 | 4.17931595 | 3.39153345 |
| TGFb1 | 0.80829529 | 2.05545484 | 1.43187506 | 0.88187498 |
| IL-3 | 1,51873155 | 1,0417141 | 1,28022282 | 0,33730228 |
| IL-6 | 1,60492919 | 5,84406016 | 3,72449468 | 2,99751826 |
| IL-8 | 1,62122753 | 2,2758561 | 1,94854182 | 0,4628923 |

**Table S21**: Fisher's exact test results of sub-compartment switches in NF-kappa target genes vs. non-NF-kappa target genes

| Switch | Odds ratio | P-value |
|---|---|---|
| NC vs. C15m | 1.33 | 0.022 |
| NC vs. C4h | 1.053 | 0.713 |
| C15m vs. C4h | 0.996 | 1 |
| NC vs. C15m vs. C4h | 1.145 | 0.299 |

**Table S22: Publicly available data used for subcompartment overlap comparison analysis**

| Biosample | Assay | Target | Dataset accession | bed narrowPeak file accession |
|---|---|---|---|---|
| Homo sapiens IMR-90 | Histone ChIP-seq | H3K9me3 | ENCSR055ZZY | ENCFF098XMT |
| Homo sapiens IMR-90 | Histone ChIP-seq | H3K27ac | ENCSR002YRE | ENCFF805GNH |
| Homo sapiens IMR-90 | Histone ChIP-seq | H3K27me3 | ENCSR431UUY | ENCFF336IXL |
| Homo sapiens IMR-90 | Histone ChIP-seq | H3K4me3 | ENCSR087PFU | ENCFF093NQC |
| Homo sapiens IMR-90 | Histone ChIP-seq | H3K9ac | ENCSR219MYH | ENCFF309ZMM |
| Homo sapiens IMR-90 | Histone ChIP-seq | H3K4me1 | ENCSR831JSP | ENCFF611UWF |
| Homo sapiens IMR-90 | TF ChIP-seq | CTCF | ENCSR000EFI | ENCFF203SRF |
| Homo sapiens IMR-90 | ATAC-seq | DNA | ENCSR200OML | ENCFF243NTP |
| Homo sapiens IMR-90 | ChIP-seq | Lamin B1 | GSE36641 | GSE36616 |
| Homo sapiens IMR-90 | Whole-Genome Tiling Array | Nucleolus-associated DNA | GSE78043 | S1 table |

**Table S23: The total number of significant inter- and intra-chromosomal interactions in each sample identified by NCHG**

| Sample | Cis (FDR = 0.01; logRatio=2.0) | Trans (FDR = 0.01; logRatio=1.5) |
|---|---|---|
| NC rep1 | 23068 | 10014 |
| NC rep2 | 16112 | 12307 |
| NC rep3 | 14279 | 20179 |
| NC consensus | 16861 | 10411 |
| C15m rep1 | 22858 | 20965 |
| C15m rep2 | 23214 | 11167 |
| C15m rep3 | 22907 | 59081 |
| C15m consensus | 21214 | 17681 |
| C4h rep1 | 19159 | 11457 |
| C4h rep2 | 4744 | 22767 |
| C4h rep3 | 20096 | 8062 |
| C4h consensus | 13419 | 11110 |
